# Genetic associations at regulatory phenotypes improve fine-mapping of causal variants for twelve immune-mediated diseases

**DOI:** 10.1101/2020.01.15.907436

**Authors:** Kousik Kundu, Alice L. Mann, Manuel Tardaguila, Stephen Watt, Hannes Ponstingl, Louella Vasquez, Nicholas W. Morrell, Oliver Stegle, Tomi Pastinen, Stephen J. Sawcer, Carl A. Anderson, Klaudia Walter, Nicole Soranzo

**Author notes:** Address for correspondence: Prof. Nicole Soranzo, Human Genetics, Wellcome Sanger Institute, Genome Campus, Hinxton, CB10 1HH.

## Abstract

The identification of causal genetic variants for common diseases improves understanding of disease biology. Here we use data from the BLUEPRINT project to identify regulatory quantitative trait loci (QTL) for three primary human immune cell types and use these to fine-map putative causal variants for twelve immune-mediated diseases. We identify 340 unique, non major histocompatibility complex (MHC) disease loci that colocalise with high (>98%) posterior probability with regulatory QTLs, and apply Bayesian frameworks to fine-map associations at each locus. We show that fine-mapping applied to regulatory QTLs yields smaller credible set sizes and higher posterior probabilities for candidate causal variants compared to disease summary statistics. We also describe a systematic under-representation of insertion/deletion (INDEL) polymorphisms in credible sets derived from publicly available disease meta-analysis when compared to QTLs based on genome-sequencing data. Overall, our findings suggest that fine-mapping applied to disease-colocalising regulatory QTLs can enhance the discovery of putative causal disease variants and provide insights into the underlying causal genes and molecular mechanisms.

## Introduction

Immune-mediated diseases (IMDs), including autoimmune diseases, are chronic health conditions that affect around up to 9.4% of the world population ^1,2^. In many chronic and debilitating immune-mediated disorders, genetic predisposition and diverse environmental factors trigger an abnormal immune response, which eventually destroy healthy tissues^3,4^. Thousands of genetic loci influencing susceptibility to different IMDs have been discovered to date by genome-wide association studies (GWAS). For a small subset of loci, genetic associations have already yielded novel insights into likely pathophysiological mechanisms underpinning disease predisposition^5,6^, for instance linking NFKB pathway genes (*NFKB1* and *TNFAIP3*) and TNF-receptor gene (*TNFR1*) to risk of inflammatory bowel disease, multiple sclerosis and ankylosing spondylitis ^6–8^. However, the causal variants and the molecular mechanisms underpinning GWAS associations remain largely unknown, hindering efforts to develop new treatments. The prioritisation of the most likely causal variants, and the identification of the putative molecular mechanisms through which causal variants act to deregulate immune pathways, are the necessary next steps to harness the power of these genetic discoveries.

Fine-mapping algorithms are used to prioritize causal variants for common complex diseases or traits by estimating the probability of a genetic variant being causal for a given phenotype, conditional on all associated variants in a given genomic region. Recently, statistical fine-mapping approaches have been used to resolve causal variants at IMD loci ^9–11^. However, many of the risk variants discovered to date are common (defined here as having minor allele frequencies [MAF] ≥ 5%), and exhibit high levels of linkage disequilibrium (LD) with other nearby variants. This limits our ability to resolve causal variants based on disease summary statistics data alone ^6,9,12^.

Common disease risk loci are preferentially found in non-coding regions of the genome, suggesting that the majority of these variants may exert their effects on disease through gene regulation^13–16^. Gene regulation traits (principally gene expression, splicing and chromatin phenotypes) provide a first readout of the activity of genetic variants in cell-defined contexts, which can be captured using QTL mapping. Gene expression and splicing QTLs (eQTLs and psiQTLs) often colocalise with disease association signals ^17–20^. Further, evidence for colocalisation extends beyond gene expression and splicing, to QTLs for DNA methylation and histone modifications (e.g., marking active enhancers and promoters), providing information complementary to eQTLs to link regulatory elements to the genes they control^20,21^. When there is a shared genetic signal, these gene regulation QTLs can be leveraged to improve the resolution of fine-mapping.

The main advantage of using QTLs for fine-mapping causal variants is that they typically exhibit larger effect sizes per variant than complex disease odds ratios^22,23^, thus achieving the same statistical power by using order of magnitude smaller numbers of individuals than GWAS-based fine-mapping studies. Further, the smaller size of these studies makes it economically feasible to derive regulatory QTLs from complete genetic maps based on whole genome sequencing (WGS), thus potentially increasing the resolution of fine-mapping compared to datasets based on sparser imputation panels.

Here we sought to quantify the value of regulatory QTLs to resolve causal variants across a range of common immune-mediated diseases, where compared to currently-available gold-standard disease datasets based on meta-analyses of many smaller studies (**Figure 1**). We show that QTL data improves resolution of causal variants compared to disease summary statistics alone, and identifies putative effector genes, cell types, and regulatory mechanisms for causal variant, thus enhancing interpretation of putative causal functional effects for disease variants.

**Figure 1:**
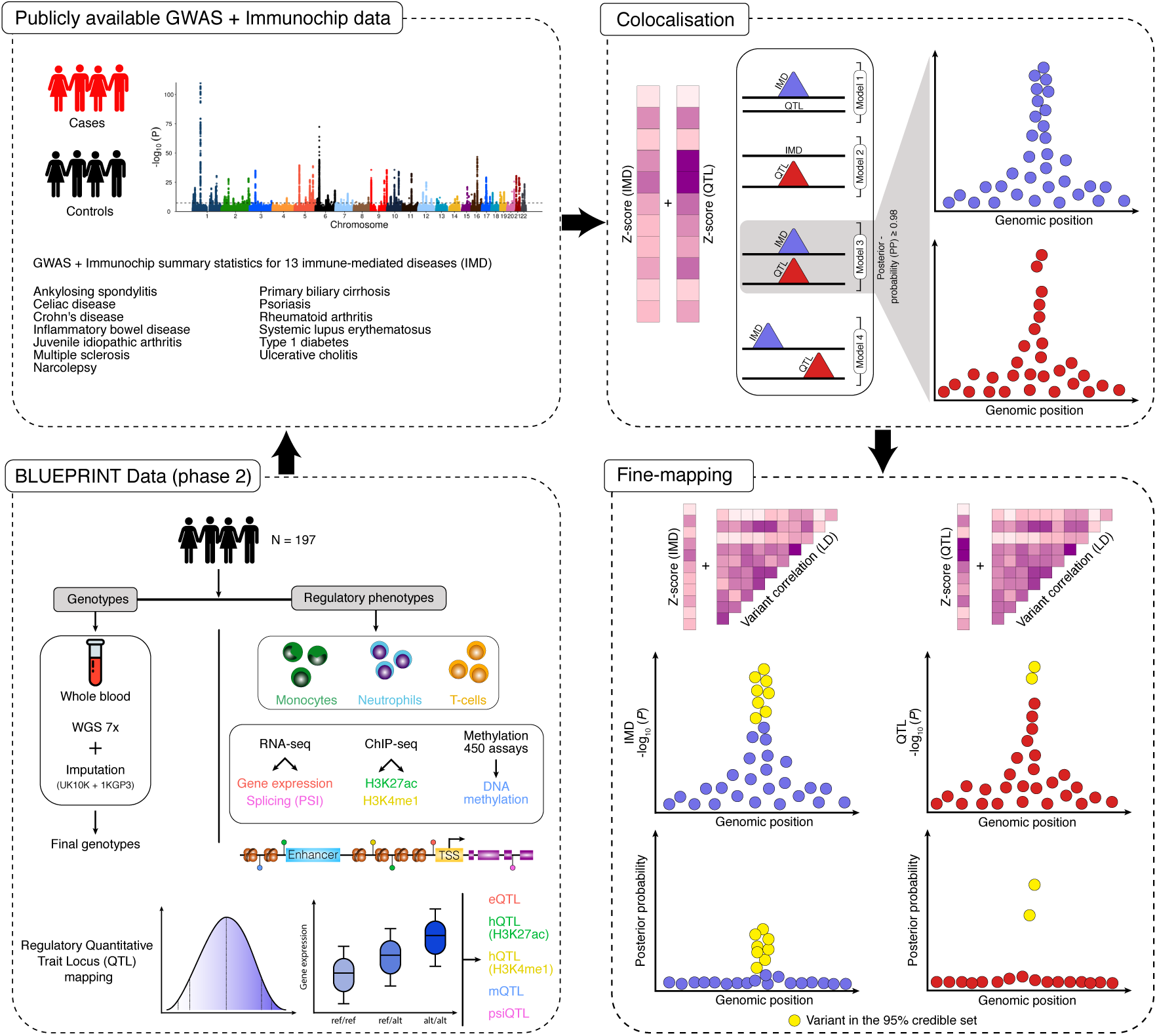
Study workflow. Figure describes the overview of study design. We first compiled publicly available IMD loci for 13 diseases and re-computed QTLs as part of the BLUEPRINT phase 2 data. We then performed statistical colocalisation on QTL and IMD loci, and used these regulatory QTLs to explain putative path from genetic variants to disease. Finally, we performed genetic fine-mapping to identify potential causal variants and also evaluated if regulatory QTLs lead to improvements in fine-mapping compared to disease summary statistics alone.

## Results

### The BLUEPRINT human variation phase 2 dataset

As part of the BLUEPRINT project, we previously generated regulatory datasets for three primary human immune cell types (i.e., CD14+ monocyte, CD16+ neutrophil, and CD4+ T-cell), including transcription, histone binding and DNA methylation phenotypes across 197 individuals drawn from a UK-based bioresource ^20^. In the Chen et al. study ^20^, conservative parameters were applied to call variants from low-read depth (7x) WGS data, at the expense of sensitivity of variant discovery. To increase sensitivity to discover potential causal variants, here we reprocessed the WGS dataset using an alternative set of parameters (99.6% and 90% truth sensitivity for single nucleotide polymorphisms (SNPs) and INDELs, respectively) and performed an additional genotype refinement step ^24^ followed by an imputation strategy using the combined UK10K and 1000 Genomes Project Phase 3 (UK10K + 1KGP3) haplotype reference panel (**Methods; Supplementary Figs. 1-2**). The resulting ‘phase 2’ dataset contains a total 9,228,816 variants (8,320,384 SNPs and 908,432 INDELs; approximately 1.4 million more SNPs and almost 10 times more INDELs when compared to the phase 1 data; **Supplementary Table 1 and Supplementary Fig. 3**). We applied linear mixed models ^25^ to recompute QTLs using the phase 2 genetic variants and different classes of regulatory features, including for gene expression levels (eQTLs), percent spliced-in (psiQTL), H3K27ac and H3K4me1 histone binding (hQTL), and DNA methylation (mQTL). We tested cis-associations in 1Mb windows centered on each feature (e.g., gene, methylation probe) in three different primary blood cell types, i.e., monocyte, neutrophil, and T-cell (**Methods**). The phase 2 dataset captures 99% of phase 1 QTL signals at 5% global False Discovery Rate (gFDR) correction, considering sentinel variants that are either the same as in phase 1, or a close proxy (*r*^2^ ≥ 0.8; **Methods; Supplementary Fig. 4**). The phase 2 data and QTL summary statistics are released through the European Genome-phenome Archive (EGA accession number: EGAD00001005192, EGAD00001005199, and EGAD00001005200).

**Table 1:**
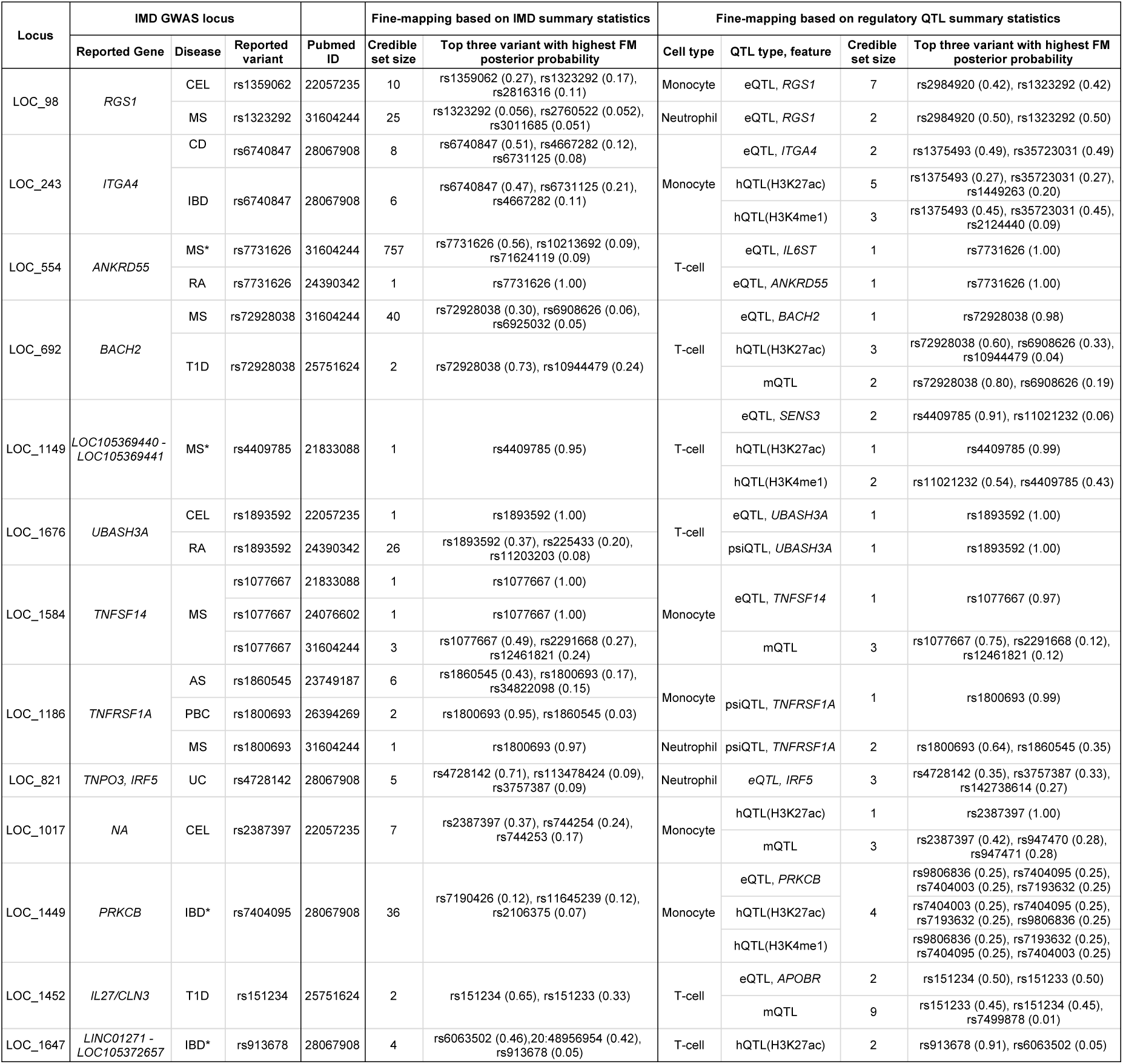
Examples of high confidence fine-mapping of IMD loci. Listed IMD loci were fine-mapped by disease summary data and confirmed by QTLs with high confidence. In most cases, IMD and QTL fine-mapping results point to the same causal variants. However, the likely causal variants yielded higher posterior probabilities (*PP*_*fm*_ ≥ 0.25) in QTL fine-mapping compared to IMD fine-mapping. Top three variants with their respective posterior probabilities (in the parenthesis) are mentioned for each locus. * denotes the locus had moderate p-value (*P* ≤ 1 × 10^−5^) but did not reach genome-wide significance p-value threshold (*P* ≤ 5 × 10^−8^) in respective IMD summary statistics.

### Colocalisation of regulatory QTLs with immune-mediated disease loci

We first sought to identify IMD loci sharing a genetic signal with the phase 2 QTL data. We retrieved publicly available GWAS summary statistics for 13 IMDs, including seven previously analysed in Chen et al. ^20^ (celiac disease [CEL], Crohn’s disease [CD], inflammatory bowel disease [IBD], multiple sclerosis [MS], rheumatoid arthritis [RA], type 1 diabetes [T1D], and ulcerative colitis [UC]) and six not previously investigated (ankylosing spondylitis [AS], juvenile idiopathic arthritis [JIA], narcolepsy [NAR], primary biliary cirrhosis [PBC], psoriasis [PSO], and systemic lupus erythematosus [SLE]) ^8,26–44^. Overall, we considered a total of 31 datasets, including 18 datasets (updated Jan 2019, **Supplementary Table 2**) generated via commercial genotype arrays, and 13 generated using bespoke-content arrays (Immunochip) ^45,46^.

To identify IMD loci that are likely to share a causal variant with one or more of our regulatory QTLs, we first systematically searched the genome-wide statistics to identify loci where the IMD sentinel SNP or a proxy (*r*^2^ ≥ 0.8; **Methods**) is also a sentinel variant in the QTL dataset. We excluded the human leukocyte antigen (HLA) region where standard SNP tagging approaches do not perform consistently well ^47^. This approach identified 5,257 LD-overlapping IMD/QTL pairs. We then used a Bayesian colocalisation method (pw-gwas ^48^) to compute prior probabilities of each regional model from the maximum log-likelihood function of all variants in the tested region. Among the 5,257 pairs, 4,819 (92%, corresponding to 340 IMD associations) had robust statistical support for colocalisation (≥ 98% posterior probability [*PP*_*coloc*_]), while 438 (8%) had robust statistical support for linkage, indicates that the signals for IMD and QTL are independent (**Methods; Figure 2a; Supplementary Figs. 5-6 and Supplementary Tables 3-4**). Colocalisation was observed across all three immune cell types at 167 (49%) of the 340 loci, across two cell types at 70 (21%) loci, and specific to one of the three cell types at 103 (30%) loci (**Figure 2b**). To explore the cell type specificity of the colocalisations across a wider set of cell types and tissues, we further compared the immune cell eQTLs to a multi-tissue eQTL panel (GTEx consortium v7; **Methods**). We tested 211 immune cell eQTLs (total 150 genes) colocalising with 132 distinct IMD loci (72 distinct disease loci). Of these, more than half (109 eQTLs for 82 genes) were only observed in our immune cell types. Approximately one quarter (48 eQTLs for 36 genes) were shown to colocalise with eQTLs in a small number (≤ 4) of additional cell types. The remaining quarter (54 eQTLs for 32 genes) were highly pleiotropic (i.e., seen in 5+ more tissues; **Supplementary Fig. 7 and Supplementary Table 5**).

**Figure 2:**
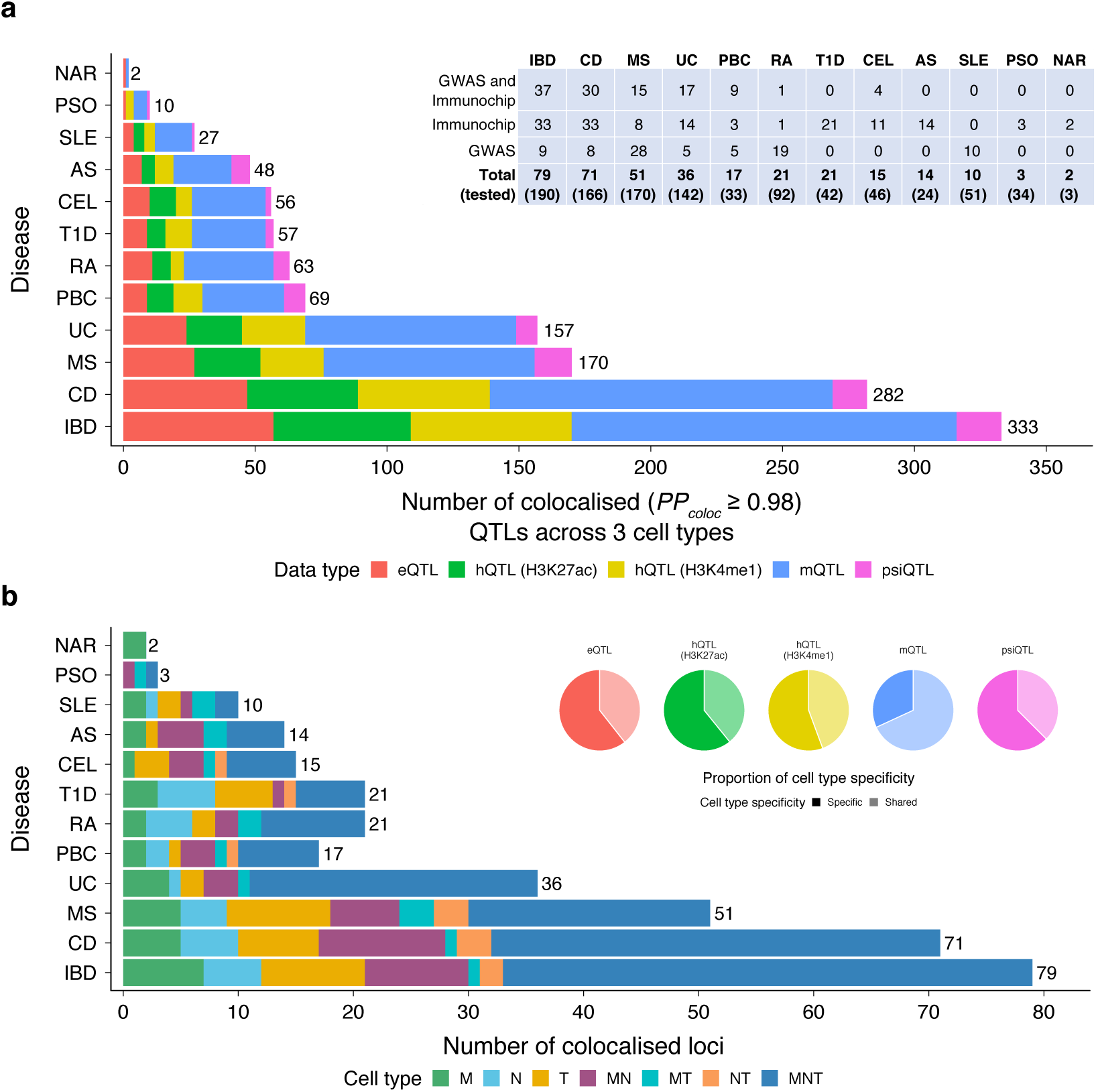
Details of IMD-QTL colocalisation. **a**, Number of IMD loci colocalised with five different QTLs across three different cell types. None of the JIA loci showed any strong colocalisation (*PP*_*coloc*_ ≥ 0.98) evidence. The table indicates the number of unique IMD loci colocalised with at least one QTL in one cell type out of the total number of loci tested (in parenthesis) along with the data sources (GWAS, Immunochip, or both) of these loci. **b**, Number of unique IMD loci colocalised with QTL and cell type information. The cell type specificity of each QTL is depicted in pi charts. This figure shows except mQTLs, all other QTLs are relatively specific to a certain cell type.

### Regulatory QTLs more accurately define disease causal variants compared to disease summary statistics

To compare fine-mapping results across disease and regulatory data, we first focused on a set of 124 loci where the density of genetic variants in the QTL and corresponding IMD summary statistics were comparable. Specifically, we required that at least 80% of the genetic variants found in each 1Mb QTL genomic interval were also found in the IMD dataset, and vice-versa. For simplicity, we also focused only on regions containing a single association signal, as inferred by stepwise conditional analysis for IMD loci or exact conditional tests based on individual-level genetic data for QTLs (**Methods**).

We carried out fine-mapping separately on QTL and IMD summary statistics, setting the number of input causal variants to one per locus (**Methods**). We used two Bayesian fine-mapping frameworks, namely FINEMAP^51^, which uses an efficient shotgun stochastic search algorithm, and CAVIARBF ^52^, which uses an exhaustive search algorithm. Both methods yielded near-identical results after parameter optimization (**Methods; Supplementary Fig. 8 and Supplementary Table 6**), so for simplicity we present here the FINEMAP results.

At each colocalised locus, we derived 95% credible sets from QTL and IMD summary statistics separately. Namely, for each fine-mapping experiment, we ranked each variant by decreasing fine-mapping posterior probability (*PP*_*fm*_), and selected the minimal set of variants that jointly accounted for ≥ 95% *PP*_*fm*_. We first compared the size of the 95% credible sets between the QTL and IMD fine-mapping (**Figure 3a**). When considering the minimal credible set out of all QTLs colocalising with each given IMD locus (i.e. spanning five traits and three cell types), QTLs yielded 2,741 variants across 124 loci, and IMDs yielded 4,902 variants (t-test *P* = 4.8 × 10^−4^; **Figure 3a**). Overall, QTLs produced smaller credible sets than IMD for 70% (87/124) of colocalising loci. Furthermore, on average the top variant in each QTL credible set achieved a higher *PP*_*fm*_ compared to that from the colocalising IMD (mean *PP*_*fm*_ = 0.40 vs 0.27, t-test *P* = 1.9 × 10^−5^; **Figure 3b**). Out of 124 colocalising loci, QTL fine-mapping resolved 14 (11%), 30 (24%) and 39 (32%) loci to credible sets of 1, 2-5, 6-20 variants, respectively, compared to 2 (2%), 14 (11%) and 43 (35%) for IMD summary statistics (**Figure 3c**). The same trend of smaller credible sets for QTLs was observed when compared to a recent fine-mapping study in IBD based on a larger sample size^9^ (**Methods; Supplementary Fig. 9 and Supplementary Table 7**).

**Figure 3:**
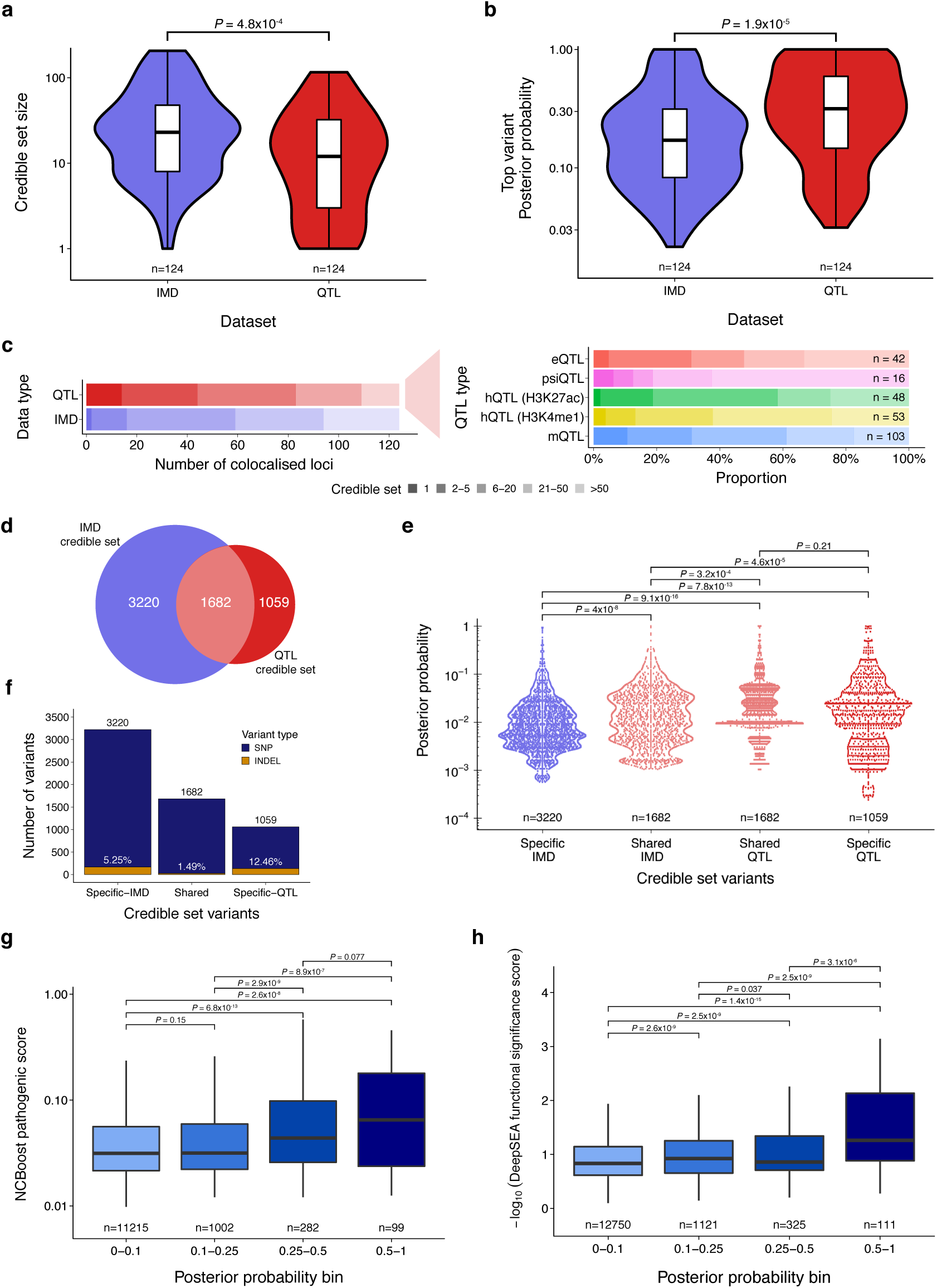
Fine-mapping of QTL and IMD loci. **a**, Credible set size comparison between QTLs and IMD loci. For QTLs, we considered the smallest credible set derived from five different QTLs in three different cell types. **b**, Comparison of the top most variant for each credible set, which achieved highest posterior probability (*PP*_*fm*_) in QTL and IMD fine-mapping. In both figures (a and b), a two sided t-test showed there is a significant difference in two credible sets. **c**, Number of variants in QTL and IMD credible sets depicted for colocalised loci. This figure indicates QTL fine-mapping achieves smaller credible sets than IMD fine-mapping. For example, 44 disease loci were fine-mapped to maximum five variants credible set where the equivalent credible set size was achievable for only 16 disease loci by IMD fine-mapping. The proportion of credible set size for each QTL is also reported. **d**, Venn diagram showing the overlapping variants in the credible sets between QTL and IMD loci. **e**, Posterior probability (*PP*_*fm*_) distribution of shared (intersection) and specific variants for QTL and IMD credible sets. A t-test is used to calculate significance (*P*) values. **f**, Proportion of variants (SNP/INDEL) in shared, specific QTL, and IMD credible sets. **g**,**h**, Pathogenic score (via NCBoost ^49^) and functional significance score (via DeepSEA^50^) distributions of variants where the credible set ≤ 50 are depicted in different posterior probability bins. Both figures indicate that the variants with higher *PP*_*fm*_ were also yielded higher pathogenic and/or functional significance score, and the differences were statistically significant (Wilcoxon test).

We next compared posterior probabilities for putative causal variants in the 95% credible sets. Overall, 1,682 variants were in both the IMD and QTL credible sets, corresponding to approximately two-thirds of all QTL and one-third of all IMD credible variants respectively (intersection set; **Figure 3d**). Within the intersection set, variants had marginally higher posterior probabilities in the QTL compared to the IMD analysis (shared QTL *PP*_*fm*_ mean = 0.042 and shared IMD *PP*_*fm*_ mean = 0.031, t-test *P* = 3.2 × 10^−4^, **Figure 3e**). Variants only included in the IMD credible sets had lower overall *PP*_*fm*_ (mean = 0.02) compared to variants in the intersection set (mean = 0.031, t-test *P* = 4 × 10^−8^; **Figure 3e**). Conversely, credible variants in the QTL-specific set had similar PPs to those in the high-confidence intersection set (mean = 0.047, t-test *P* = 0.21; **Figure 3e**; **Supplementary Fig. 10**). Importantly, 12.5% of the candidate causal variants in the QTL-specific set were INDELs, compared to 1.49% in the intersection set and 5.25% in the IMD-specific set (**Figure 3f**). This likely reflects the systematic removal of INDELs in many published studies, and suggests that this category of putative causal variants may be systematically omitted in fine-mapping studies based on published disease summary statistics.

To predict pathogenicity of the variants in QTL data where the credible set size ≤ 50, we used a supervised learning based method, NCBoost ^49^, which uses a number of features relevant to natural selection along with interspecies conservation scores derived from different evolutionary timescales. When annotating variants in QTL credible sets using NCBoost, we found that variants in the highest fine-mapping posterior probability group (*PP*_*fm*_ = 0.5 − 1) had greater overall pathogenic scores (median = 0.065) compared to variants with intermediate (*PP*_*fm*_ = 0.25 − 0.5, median = 0.044, Wilcoxon test *P* = 0.077), low (*PP*_*fm*_ = 0.1−0.25, median = 0.032, *P* = 8.9×10^−7^) or very low (*PP*_*fm*_ = 0−0.1, median = 0.031, *P* = 2.6 × 10^−8^) posterior probability group (**Figure 3g**). Furthermore, we used a deep-learning based method, DeepSEA^50^, which predicts variant effects based on various chromatin features (e.g., transcription factor binding, histone marks, and DNase I hypersensitive sites) in multiple human cell types. DeepSEA combines these chromatin features along with evolutionary conservation to measure a functional significance score for each variant. Variants in the highest posterior probability group (*PP*_*fm*_ = 0.5 − 1) were assessed as more functionally significant (median of significance score = 0.055) compared to other groups (*PP*_*fm*_ = 0.25 − 0.5, median = 0.138, Wilcoxon test *P* = 3.1 × 10^−6^; *PP*_*fm*_ = 0.1 − 0.25, median = 0.119, *P* = 2.5 × 10^−9^; *PP*_*fm*_ = 0 − 0.1, median = 0.147, *P* = 1.4 × 10^−15^; **Figure 3h**). Together these analyses suggest our identified causal variants are more likely to be functional and have larger pathogenic effects. The pathogenic and functional significance scores for variants in QTL credible sets are reported in **Supplementary Table 8**.

### A fine mapping resource for immune-mediated diseases

To generate a fine-mapping resource for IMDs, we extended the fine-mapping analysis to the complete set of 340 colocalising IMD loci, thus removing the requirement for the QTL and IMD datasets to have a similar density of markers. Overall, we investigated 4,819 QTL-IMD pairs that colocalised with at least one QTL from at least one cell type at gFDR ≤ 5%. Out of the 340 loci, QTL fine-mapping resolved 36 (11%), 74 (22%), and 112 (33%) loci to 95% credible sets of 1, 2 − 5, 6 − 20 variants, respectively (**Supplementary Table 6**), adding to our understanding of causal variants in IMDs.

Out of all 18 loci that we considered well resolved using QTL data (credible set size ≤ 5) but not using IMD summary data (credible set size ≥ 20), there were seven loci implicating either one eQTL and/or psiQTL. One of the examples was the *BACH2* (transcription regulator protein) locus. The A allele at the intronic variant rs72928038 increases risk for MS^44^ and T1D ^41^, and the disease associations were confidently colocalised with multiple QTLs (eQTL: decreasing gene expression, psiQTL, hQTL(H3K27ac) and mQTL; *PP*_*coloc*_ ≥ 0.98; **Figure 4a; Supplementary Fig. 11**), predominantly in naive CD4+ T-cells. The sentinel variant, rs72928038, lies in the intron closest to the transcriptional start site of *BACH2*. Using T-cell Promoter Capture Hi-C (PCHi-C) data^53^, we observed a significant chromatin interaction between rs72928038 and the *BACH2* promoter (Chicago score = 9.28), supporting a spatial contact between the two (**Figure 4b**). Fine-mapping of the locus using QTL data yielded smaller credible sets for expression (n=1 variant; rs72928038; *PP*_*fm*_ = 0.98), methylation (n=2), splicing (n=3), and H3K27ac (n=3) compared to fine-mapping using MS summary statistics^44^ (n=40 variants; **Table 1; Figure 4c**), where rs72928038 achieved the highest *PP*_*fm*_ of 0.3. In T1D fine-mapping, rs72928038 was also the most likely causal variant (*PP*_*fm*_ = 0.73), although in this case the summary data was very sparse and the majority of the variants in the region were not tested^41^ (**Figure 4c**). For psiQTL, there were three variants in high LD (*r*^2^ ≥ 0.84) in the credible sets: rs6908626 (*PP*_*fm*_ = 0.5), rs72928038 (*PP*_*fm*_ = 0.24) and rs10944479 (*PP*_*fm*_ = 0.24). The psiQTLs was predicted to elicit a significant increase in the relative contribution of a processed transcript to the total transcriptional output of *BACH2* in homozygous carriers of the effect alleles (ENST00000481150; *P* = 6 × 10^−4^ for rs10944479, *P* = 7 × 10^−3^ for rs6908626, *P* = 1 × 10^−3^ for rs72928038; 5% in hom REF vs 10% in hom ALT). This suggests that the effect of the downregulation of the eQTL is exacerbated by the splice QTL due to an increased contribution of non-coding isoforms (**Figure 4d**). Another example was rs1893592, located three bases downstream from the tenth exon of *UBASH3A*, and associated with RA, CEL, and PSC ^27,38,56^. *UBASH3A* encodes a protein belonging to the T-cell ubiquitin ligand family that negatively regulates T-cell signalling. The A risk allele was associated with decreased gene expression and increased percent-splice-in (PSI) in our study, supporting previous evidence ^57,58^. The locus could be fine-mapped to a single variant credible set (rs1893592, *PP*_*fm*_ = 1) using either eQTL or psiQTL, while IMD fine-mapping for RA ^38^ yielded a credible set of 26 variants, where *PP*_*fm*_ for rs1893592 = 0.37 (**Table 1; Supplementary Fig. 12**).

**Figure 4:**
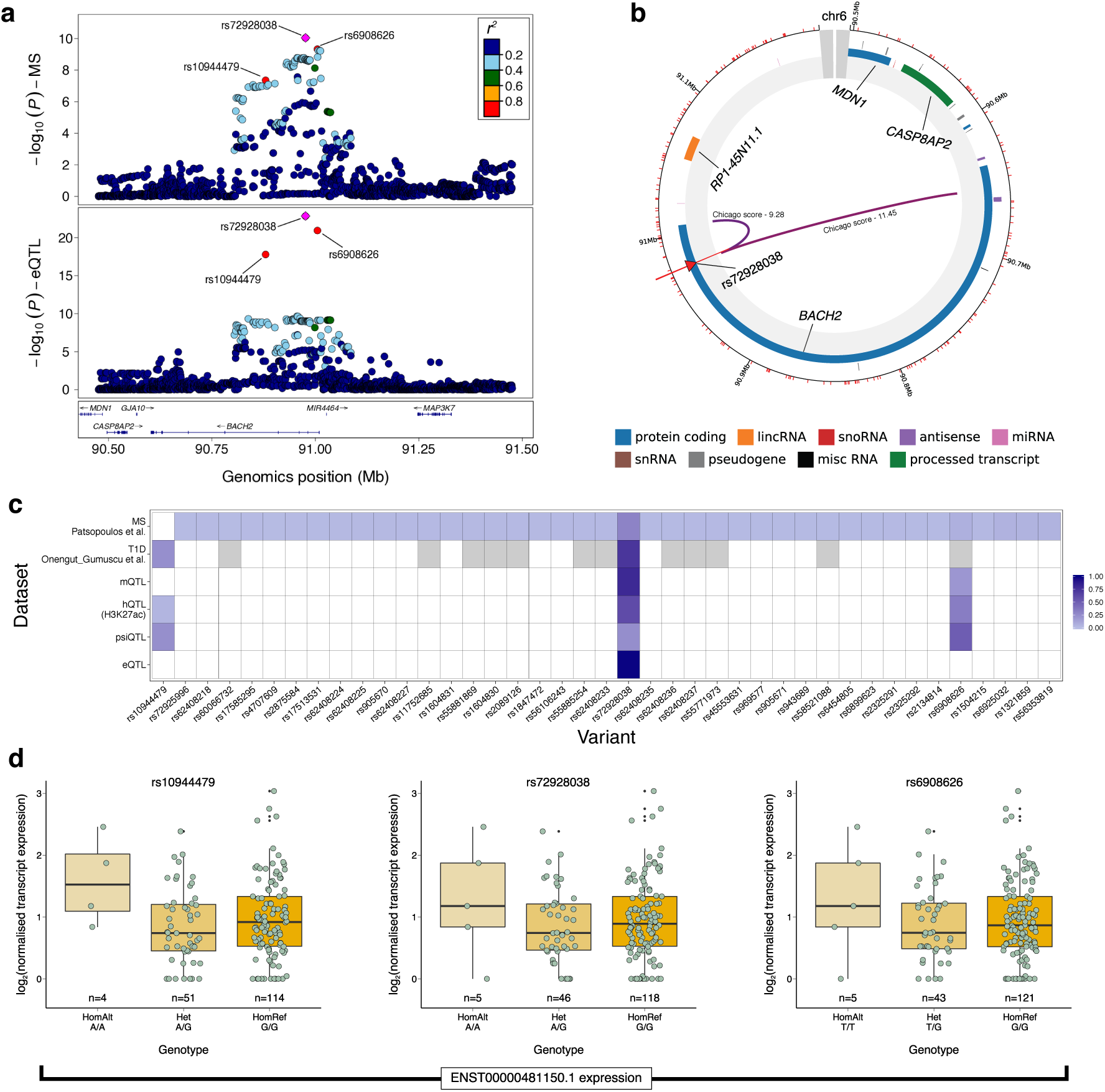
Fine-mapping of the BACH2 locus in CD4+ T-cells. **a**, The locuszoom plot of the *BACH2* locus using eQTL and MS data with 500kb flanking region surrounding the sentinel SNP rs72928038. Pairwise LD (*r*^2^) was calculated using BLUEPRINT data. The rs72928038 is an intronic variant of *BACH2* and also known to be a risk allele (A) for MS ^44^ and T1D ^41^. The locus was strongly colocalised (*PP*_*coloc*_ ≥ 0.98) between IMDs (MS and T1D) and multiple QTLs (i.e., eQTL, psiQTL, hQTL(H3K27ac) and mQTL) in naive T-cells. **b**, The sentinel variant, rs72928038, resides near the Transcriptional Start Site (TSS) of *BACH2*. A significant chromatin interaction between rs72928038 and promoter of the *BACH2* (Chicago score of 9.28) was observed in naive CD4+ T-cell using PCHi-C data ^53^, which shows a strong regulatory effect. The figure was generated by using CHiCP web server ^54^. **c**, Heatmap shows posterior probability of the fine-mapped variants (*PP*_*fm*_) in the respective credible sets (colour intensity: PP_*fm smallest*_ - light blue to PP_*fm largest*_ - deep blue). White colour indicates the variants were not part of the respective credible sets. Due to lack of variant density in T1D summary statistics, many variants were not tested for fine-mapping analysis (grey). The locus could not be fine-mapped confidently with MS summary statistics, yielded a credible set of 40 variants (rs72928038 being the top variant; *PP*_*fm*_ = 0.3), whereas a single variant (rs72928038; *PP*_*fm*_ = 0.98) credible set was achieved by using eQTL data. The rs72928038 appeared to be the most likely causal variant in all other data sets as well, except psiQTL (second highest; *PP*_*fm*_ = 0.24). **d**, Transcript expression levels for psiQTLs. The two variants rs10944479 (*r*^2^ = 0.87 with rs72928038) and rs6908626 (*r*^2^ = 0.95 with rs72928038) were part of the psiQTL and hQTL(H3K27ac) credible sets (c). These two variants along with the sentinel variant were characterized by an increased exon-skipping of *BACH2* exon 8 (coordinates - 6:90,798,680−90,798,772). The psiQTLs indicate a significant increase in the relative contribution of a processed transcript to the total transcriptional output of the *BACH2*.

There were several other examples where our QTL fine-mapping showed greater resolution to identify potential causal variants than IMD summary statistics (**Table 1**). The *ITGA4* locus encodes Integrin Subunit Alpha 4, and was recently associated with IBD ^29^. The monoclonal antibody Vedolizumab specifically binds the *α*4*β*7 integrin dimer formed by *ITGA4* and *ITGB7*, reducing gastrointestinal inflammation in IBD^59,60^. Fine-mapping of IBD associations at this locus using IBD GWAS summary statistics (**Methods**) yielded 11 variants, of which rs6740847 had the highest *PP*_*fm*_ (0.21). Fine-mapping of the same locus using monocyte QTLs yielded smaller credible sets for expression (n=2 variants), H3K4me1 (n=3) and H3K27ac (n=5) QTLs, all nested within the disease credible set (**Figure 5**). The 95% eQTL credible set consisted of one SNP and one INDEL variants in complete LD and with identical *PP*_*fm*_ (rs1375493 G/A and rs35723031 G/GT, PP=0.49; *r*^2^=1; **Figure 5a-b; Supplementary Fig. 13**). However, the INDEL (rs35723031) was not included in any published IBD meta-analysis data ^28,29^ (**Supplementary Fig. 13c**). Analysis of transcription factor binding data showed that in monocytes the rs1375493 variant maps directly to a binding peak of the haematopoietic master regulator C/EBP*β* (**Figure 5c**). Analysis of regulatory scores using DeepSEA^50^ showed that the QTL lead SNP rs1375493 achieved highest functional significant score among all 11 variants at this locus, and interestingly the chromatin feature effect was most significant for H3K27ac in untreated monocytes (*E* − *value* = 1.82 × 10^−4^; **Supplementary Fig. 14**). As the colocalisation evidence of *ITGA4* locus with stimulated monocytes was reported previously ^29^, we analysed this locus in stimulated monocytes from two different studies ^61,62^. Both studies showed that the lead SNP rs1375493 had stronger association in all stimulated conditions when compared to the IBD lead variant rs6740847 (INDEL: rs35723031 was not tested in both studies^61,62^; **Supplementary Fig. 15**). These and other examples (*TNFSF14* and *SESN3* loci for MS, *RGS1* for MS and CEL, *TNFRSF1A* for MS, PBC and AS, and *APOBR* for T1D; **Supplementary Fig. 16-20**) demonstrate the power of regulatory QTLs for identifying causal variants, and informs downstream disease mechanism studies. We also reported loci where IMD and QTL (mainly eQTL) fine-mapping indicate same causal variants (**Table 1**). Remarkably, as in the *ITGA4* case above, INDELs accounted for over 12% of QTL-specific credible variants, for instance and were contained in the high-probability (*PP*_*fm*_ ≥ 0.25) credible sets at 22 other loci (e.g., *IRF5* for UC, *PARK7* for IBD and *SH2B3* for AS; **Supplementary Table 9**). This illustrates the value of using INDEL imputation reference panel or genome-wide sequencing data to achieve a more comprehensive evaluation of potential causal genetic variants in fine-mapping studies.

**Figure 5:**
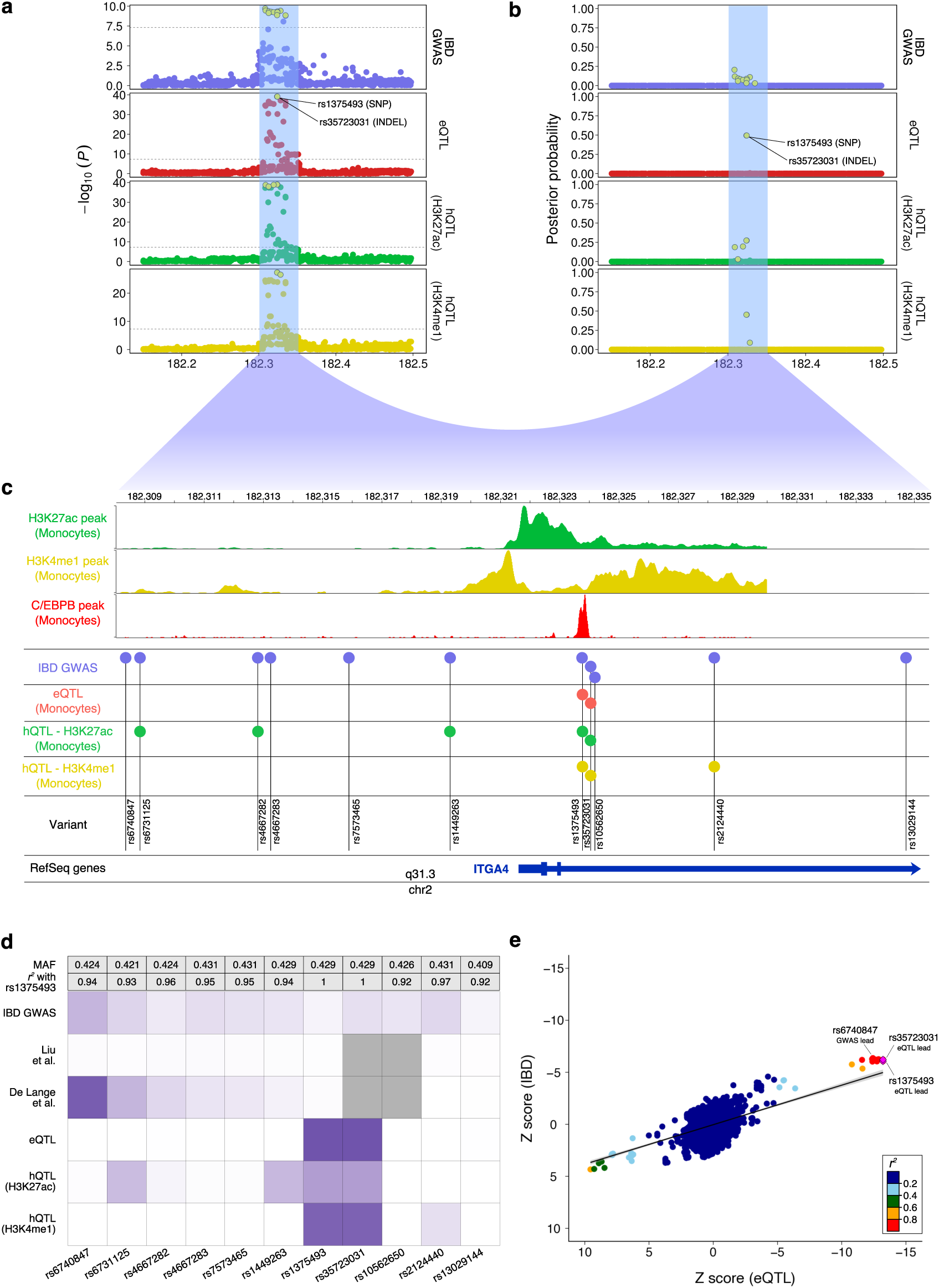
Fine-mapping of the ITGA4 locus in monocyte. **a**,**b**, Fine-mapping of IBD associated *ITGA4* locus using different QTL in monocyte and IBD GWAS summary statistics (**Methods**). Potential causal variants (95% credible set) for respective data sets (IBD GWAS: 11, eQTL: 2, H3K27ac: 5, and H3K4me1: 3) are highlighted by yellow. Among the QTL fine-mapping, eQTL provides smallest credible set consisting only two variants; one SNP (rs1375493) and one INDEL (rs35723031) falling in the second intron of the gene. **c**, Genomic plot of the *ITGA4* region where H3K27ac, H3K4me1, and C/EBP*β* (derived from^55^) peaks are depicted along with the genomic positions of all the variants in 95% credible set derived from different data sets. Most of the QTL variants fall where most chromatin activities were observed. Interestingly, one of the two eQTL variants (rs1375493) that also achieved highest posterior probability (*PP*_*fm*_) in all QTL data was observed to be residing within the C/EBP*β* peak, which provides more confidence that this variant might play a role in the gene regulation through promoter interaction, however this requires further experimental validation. On the other hand IBD GWAS lead variant (rs6740847; *PP*_*fm*_ = 0.21) falls 13kb upstream of the gene and the region does not show any chromatin activities. **d**, Posterior probability heatmap of all the 11 variants derived from IBD GWAS and QTL data sets. In addition to IBD GWAS data, we used two additional IBD meta analysis data (Liu et al. ^28^ and De Lange et al. ^29^). All the likely causal variants in QTL credible set were subset of IBD GWAS credible set. Both eQTL fine-mapped variants were in absolute LD (*r*^2^ = 1) and therefore achieved exactly the same posterior probability (*PP*_*fm*_ = 0.49), which was higher than any other variant in the region. Minor allele frequency and pairwise LD values (with QTL lead: rs1375493) for each of the 11 variants are mentioned. Note that the IBD meta-analysis datasets do not contain INDELs (rs35723031 and rs10562650). **e**, Z-scores derived from eQTL and IBD GWAS for the locus are plotted against each other with LD information. This plot confirms the colocalisation was meaningful, indicating both are the same signal and the variants effects are in the same direction.

There were also examples of variants regulating multiple genes. The intronic variant rs7731626 mapping to *ANKRD55* was previously reported as risk alleles for RA ^38^ and MS ^44^. The protective allele rs7731626 (A) is associated with decreased expression of *ANKRD55*, and with decreased expression of *IL6ST* in CD4+ T-cells. The *IL6ST* gene encodes Interleukin 6 Signal Transducer, a protein that allows signal transduction of the IL6 pathway^63^. Previously, Chun et al ^64^ reported an association between rs71624119 and *ANKRD55* expression but not with *IL6ST* in CD4+ T-cells. The rs71624119 variant is in moderate LD (*r*^2^ = 0.53) with the RA and MS risk allele rs7731626, and has weaker support in our analysis (*IL6ST* eQTL: rs71624119 *P* = 3.5 × 10^−10^, rs7731626 *P* = 2.3 × 10^−13^ in T-cells). The *IL6ST* eQTL is further supported by PCHi-C data that shows an interaction between the variant and the promoter of *IL6ST* specifically in naive CD4+ T-cells. Overall, these findings support the IL6 signaling pathway as a druggable target for autoimmune diseases (**Supplementary Fig. 21**).

## Discussion

The main aim of this study was to assess the utility and resolution of fine-mapping methods applied to molecular QTL datasets, when compared to the current gold standard based on disease GWAS meta-analyses. Fine-mapping is most robust in settings where statistical power is high, where the catalog of genetic variants is complete, where all the genetic variants are perfectly genotyped, and where LD can be directly estimated from the study sample ^6,12,65^. Efforts to build such datasets by sequencing the genomes of hundreds of thousands of cases and controls are ongoing. However, for the majority of human diseases we are still a long way away from being able to generate genome sequencing datasets of sizes comparable to current imputation-based GWAS studies, which remain the most viable approach for fine-mapping in most diseases and traits.

We fine-mapped 340 IMD association loci across 12 diseases by using five different regulatory QTL datasets profiled in three primary cell types and nearly 200 individuals. Our analysis showed that fine-mapping based on regulatory QTLs in less than 200 people yields smaller average credible sets compared to identical approaches based on disease summary statistics of hundreds of thousands of cases and controls. A main advantage of regulatory QTLs is that, owing to their average large effect sizes, they require order-of-magnitude fewer individuals to detect associations compared to disease endpoints. This makes it cost-effective to use WGS to derive a near-complete representation of genetic variants. Conversely, common imputation reference panels are by definition sparser than WGS datasets. Further, despite attempts to standardise preprocessing and overall quality metrics in meta-analyses, heterogeneity may arise from subtle differences in imputation strategy or post-GWAS filtering approaches, which may for instance lead to systematic under-representation of particular classes of variants (e.g., INDELs) in different studies ^6^. As shown in this study, incomplete representation of genetic variants in disease summary statistics leads to the systematic exclusion of high-probability causal variants. For instance, nearly 25% of the high-confidence credible set variants (*PP*_*fm*_ ≥ 0.25) identified in the regulatory QTL data were not represented in the IMD GWAS datasets, including importantly 8% of INDELs. These results highlight the importance of developing highly-complete genome sequencing datasets for the purpose of fine-mapping. The increasing size of whole-exome and whole-genome sequencing datasets for disease discovery will ultimately ameliorate this concern. In the meanwhile, regulatory QTLs provide a cost-effective alternative to this approach, reducing by orders of magnitude the number of individuals required for fine-mapping.

A second main advantage of regulatory QTLs is that they provide a more direct interpretation of biological mechanisms underlying disease variants, accelerating downstream functional validation efforts. Further, the parallel profiling of multiple levels of regulatory annotations in multiple cell types enhances the biological and contextual interpretation of causal effects, including inference on the identity of putative causal cell types or the likely location of regulatory elements. A main caveat of this approach is that colocalisation does not allow us to discriminate cases where there is a causal relationship between the QTL and IMD variants, from those where variants may have shared but independent (‘pleiotropic’) effects on both sets of traits^48^. Furthermore, causal effects may be driven by other unmeasured cell populations, and thus colocalisation approaches alone are insufficient to conclusively pinpoint the precise cellular context(s) in which many disease-associated variants may exert their causal effect. This may not be a concern when QTLs display the same patterns of association between cells and tissues, as in the *ITGA4* example where the patterns of association in resting monocytes were replicated in monocytes exposed to a variety of different stimuli. Ultimately, however, the creation of cell- and context-resolved QTLs for a large number of biological domains (e.g., cellular, developmental, stimulus-dependent), coupled with deep experimental validation, will provide the necessary frameworks to correctly interpret the effect of each disease-associated variant. Finally, many IMDs affect ethnicities differentially ^1^. At the moment, QTL datasets target predominantly European-ancestry populations. Extension of analyses of regulatory variation in more representative sets of human populations will greatly enhance our efforts to interpret genetic associations in the context of the full spectrum of human population variation.

## Methods

### BLUEPRINT phase 2 data

We created a new phase 2 variant call set from low-read depth BLUEPRINT WGS dataset ^20^ as follows.

#### Variant quality score recalibration (VQSR)

After calling all the raw variants of 200 samples using samtools/bcftools (see Chen et al. ^20^ for details), GATK (v3.4) VQSR was applied separately to SNPs and INDELs on each chromosome to derive a variant quality score log odds (VQSLOD) for each variant. We set the VQSLOD threshold -1.0707 for SNPs (99.6% truth sensitivity) and 2.1094 for INDELs (90% truth sensitivity; **Supplementary Fig. 2**). We filtered out all the variants that did not pass the VQSLOD thresholds. Additionally, we removed the SNPs and INDELs that were found within three and ten base pairs of an INDEL, respectively.

#### Variant normalization

The VCF files were normalized using the vt (v0.5) software^66^, which includes two steps: (i) parsimony, where all the variants are represented in as few nucleotides as possible and (ii) left-alignment, where the start position of the variants are shifted towards the left to align to the reference genome (GRCh37).

#### Genotype refinement and imputation of variant calls to WGS reference datasets

In order to improve the accuracy of individual genotype calls, we applied a genotype refinement step on each chromosome separately using BEAGLE 4.1 (21Jan17.6cc)^24,67^, setting the modelscale parameter to 2.0 to increase the speed of the process without loss of accuracy. To infer unobserved genotypes at non-genotyped common variants, we performed a genotype imputation process using the combined UK10K and 1KGP3 WGS reference panel. This panel consists of a total 6,285 samples (3,781 UK10K and 2,504 1KGP3) and 87,558,135 bi-allelic sites. Note we did not use the more recent, larger Haplotype Reference Consortium (HRC) reference panel since (i) we only considered associations driven by common variants (MAF ≥ 5%); and (ii) we chose to maximise inclusion of INDELs, by imputing with sequence-based reference panel. Imputation and phasing were carried out by BEAGLE 4.1 (21Jan17.6cc) with default settings^24,67^. Finally, to increase the likelihood of sites being true, we removed all the variants that were specific to our dataset, i.e., not present in the reference panel.

#### Additional variant filtering

To generate the final variant set, we retained only bi-allelic variants with the following characteristics: (i) Allelic R-Squared (AR2) ≥ 0.8; (ii) Hardy-Weinberg equilibrium (HWE) *P* ≥ 1 × 10^−3^, and (iii) allele count (AC) > 4. Our final variant call set contained a total of 9,228,816 sites, including 8,320,384 SNPs and 908,432 INDELs (**Supplementary Table 1**). More detailed statistics were generated using bcftools stats (**Supplementary Fig. 1)**.

#### Quantitative trait locus (QTL) mapping

We followed an identical strategy to Chen et al. ^20^ to test for associations of phase 2 variant calls with regulatory phenotypes. Briefly, we remapped cis-acting QTLs for five different regulatory traits: (i) gene expression, (ii) percent spliced-in (PSI), (iii) H3K27ac histone modifications, (iv) H3K4me1 histone modifications, and (v) methylation levels in three different primary cell types: (i) monocyte, (ii) neutrophil, and (iii) T-cell. We considered all the genetic variants mapping to within a 1 Mb region flanking either side of each tested feature (e.g., gene body). The cis-QTL mapping was carried out by applying linear mixed models using the Limix software package ^25^. Here we tested the association between genetic variants with aforementioned five different regulatory traits. A random effect term was included for accounting polygenic signal and sample relatedness. To control batch effects, we corrected 10 PEER factors (*K* = 10) and applied quantile normalization across donors^68^. Summary tables were generated for each trait containing all summary information for each association, including p-value and effect size (beta). All the effect sizes were aligned with the alternative allele (GRCh37) of a variant.

#### Multiple testing corrections

Multiple hypothesis testing correction for cis-QTLs was carried out using EigenMT ^69^, which estimates the number of effective tests for a trait (e.g., gene expression) by considering the LD relationships among the tested variants. This process is computationally efficient, achieving accuracy comparable to permutation methods, whilst being not as conservative as a Bonferroni correction method. Statistical significance was calculated using the Q-value ^70^, which adjusts the obtained EigenMT p-values across the traits. We considered as significant all QTLs surpassing a gFDR of 0.05. Total number of QTLs along with the proportion of SNPs and INDELs for each regulatory phenotypes are mentioned in **Supplementary Table 10**.

### Curation of IMD summary statistics

#### Compilation of publicly available IMD data

We retrieved a total 28 summary statistics datasets covering 13 different IMDs from different sources (**Supplementary Table 2**). Of these 28 datasets, 15 were based on SNP genotypes imputed to different imputation panels (“GWAS”, 8 diseases). The remaining 13 were based on the Immunochip array (12 diseases). For seven diseases, both GWAS and Immunochip data were available. For each disease, we had access to summary statistics generated from up to four independent studies. We created unified formats for all summary data to account for differences in the information provided between summary statistics and to ensure consistency. We also retrieved a list of genome-wide significant loci from either Immunobase (https://www.immunobase.org/) or the GWAS catalog (v1.0.1-e89 r2017-06-19; https://www.ebi.ac.uk/gwas/), supplemented by manual curation of published literature. Other than for the exception described below, no individual-level genotype data was available for these studies.

We excluded declared IMD GWAS loci from our analyses that did not reach genome-wide significance (*P* ≤ 5 × 10^−8^) in the corresponding summary statistics available to us. In the majority of cases, these loci reached genome-wide significance in a multi-stage discovery + replication cohort, but where we had access to summary statistics only for the discovery-phase data, the variants were not genome-wide significant.

#### Association testing and variant filtering for IBD data (IBD GWAS)

We obtained individual-level genotype data for 18,344 study participants with for CD, IBD, and UC, and matched controls. Briefly, genotypes were derived using the Illumina HumanCoreExome v12 array, where cases were genotyped on version 12.1 and controls were genotyped on version 12.0. Genotypes were then imputed using an IBD-enhanced reference panel consisting of WGS data from the UK10K and 1KGP3 projects, and other sequenced IBD samples. The final dataset consisted of 4,264 CD, 8,860 IBD, and 4,072 UC cases, respectively, and 9,484 controls. We then applied filters to achieve the final set of variants with (i) INFO ≥ 0.4 and (ii) MAF (case + control) ≥ 0.001. The final sets of data contain almost 19 millions variants for each disease, including 3,257, 3,614, 2,150 genome-wide significant (*P* ≤ 5 × 10^−8^) variants in the non-MHC region for CD, IBD, and UC, respectively. We performed a case-control GWAS similar to a previous study ^29^ using SNPTEST v2.5.2^71^, using an additive frequentist test with score method.

### Locus definition

We used a set of a total 1,703 independent LD intervals for European population derived from LDetect ^72^ (https://bitbucket.org/nygcresearch/ldetect-data). These unified loci are approximately independent LD blocks in the human genome ^72^. We used these loci to compile a set of unique and independent loci across all diseases.

### Conditional analysis

To investigate the presence of multiple independent causal variants in a locus, we performed a conditional analysis on colocalised loci for QTL and IMD loci separately. We used GCTA-COJO: conditional and joint analysis to perform the conditional analysis ^73,74^. For each colocalised locus, we first performed single-SNP association analysis conditioning on the lead (sentinel) variant of the locus. After the first round of analysis, where at least one variant remained significant (conditional *P* ≤ 5 × 10^−8^ for IMD and *P* ≤ 1 × 10^−5^ for QTL), we repeated the process on the conditional summary statistics and conditioning on a variant set, which consists of the new variant and the original lead variant. We repeat the process until no genome-wide significant conditional p-value was observed. To estimate the LD between variants, we used the UK10K + 1KGP3 dataset as a reference for IMD, as individual-level data was unavailable. However, for QTL, where we had access to individual-level data, we used the −−cojo-actual-geno option.

### Overlapping and colocalisation of QTL and IMD loci

In our previous study, we observed a significant enrichment of immune related disease associated loci (*P* ≤ 1 × 10^−5^) with all types of regulatory information^20^. We used colocalisation analysis to assess whether IMD and QTL loci mapping to the same genomic interval had high probability of sharing the same genetic signal. To reduce the number of pairwise comparisons tested, we first selected IMD-QTL locus pairs where the sentinel IMD variant (from the GWAS catalog and/or Immunobase) was also either the most associated QTL variant (gFDR ≤ 0.05), or a highly associated proxy variant (defined by the values of the LD metric *r*^2^ ≥ 0.8). For this purpose, LD information was either calculated from the BLUEPRINT WGS data using PLINK (v1.9) ^75^, or retrieved from the UK10K + 1KGP3 data when the variant was not present in the BLUEPRINT WGS panel.

We then applied gwas-pw ^48^, which assigns loci to four possible models: models 1 and 2 provide evidence of a single variant association in either of the two summary statistics applied (i.e., regulatory QTL or IMD GWAS); model 3 supports the presence of a single genetic variant associated with both the regulatory and IMD traits (“colocalisation”); and model 4 provides evidence of independent effects between the IMD GWAS and regulatory QTL, indicative of two independent genetic associated variants (“linkage”). Although there are several Bayesian colocalisation methods ^48,64,76^ available, we used gwas-pw because instead of user assigned prior, the method computes prior probabilities of each of the four models from all the variants in the tested region by using the maximum log-likelihood function. In our analysis, the prior model parameter were estimated per 1 Mb genomic interval to avoid the risk of including multiple overlapping QTL testing regions. All the models provide posterior probability for association against the null model (i.e., no association). Under each model, the method calculates the posterior probability for all variants in a genomic window, where all the variants have the equal prior probability to be causal. The final posterior probability of a given genetic locus is the sum of the integral posterior probabilities of all the variants in the locus.

All four models were applied to each region. For colocalisation test, as we preselected regions based on proxy overlapping (*r*^2^ ≥ 0.8), the higher posterior probabilities were seen for either model 3 (colocalisation) or model 4 (linkage). For declaring a region as a colocalised locus, we draw the cut-off from posterior probability distribution and applied *PP*_*coloc*_ ≥ 0.98 as a cut-off for model 3 (**Supplementary Fig. 5a**). The gwas-pw calculated different priors for colocalised and non-colocalised loci (**Supplementary Fig. 5b-d**). We excluded the HLA (*chr*6: 20, 000, 000−40, 000, 000) due to the extremely complex LD structure. Overall, a total 11,458 IMD-QTL overlapping regions were tested for colocalisation, including the reported IMD loci that did not reach genome-wide significant threshold (**Supplementary Table 4**). We note here some of the caveats of colocalisation methods: (i) they consider only one causal variant in the tested region or locus, (ii) they do not allow inference of whether a “causal” or rather a “pleiotropic” relationship exists between two traits; and (iii) they have limited power where two causal variants in high LD are independently associated with each trait.

### Overlapping with the GTEx Consortium dataset

To further investigate cell type specificity of the colocalised loci, we overlapped our eQTLs with the 47 multi-tissue eQTL data from GTEx consortium (v7) ^15^. Since our eQTLs are from blood cells, we removed “Whole Blood” from our analysis, which is expected to yield substantial overlap. We systematically searched for rest of the GTEx eQTLs where the sentinel variant or a LD-proxy (*r*^2^ ≥ 0.8) were most significant in the BLUEPRINT eQTL dataset at a gFDR level of 5% (**Supplementary Table 5**).

### Fine-mapping of colocalised loci

To identify high confidence putative causal variants at each locus, we performed genetic fine-mapping on QTL and IMD colocalised loci using two state-of-the-art methods: (i) FINEMAP ^51^ and (ii) CAVIARBF ^52^. Both methods are based on a Bayesian framework, although different computational algorithms are used in these two methods. The FINEMAP method uses Shotgun Stochastic Search (SSS) algorithm and it is much faster than CAVIARBF method, which is based on an exhaustive search algorithm. For each locus, the fine-mapping outcome contains the Bayes factor and posterior probability of each variant being causal for the association. Fine-mapping methods model the LD structure and the strength of the associations (Z-score) in a locus to identify likely causal variants. Since most of the publicly available GWAS do not provide access to individual-level genotype data, typically fine-mapping efforts rely on common haplotype reference panels for LD estimation. However, subtle differences in LD structure between the test and reference population can lead to inaccurate and/or suboptimal fine-mapping, particularly for loci with multiple independent association signals ^65^. Here we carried out genetic fine-mapping under the assumption of a single causal variant in each locus, removing loci with evidence of multiple independent associations from the conditional analysis. Additionally, we only considered variants (QTL and IMD) with MAF ≥ 5% for fine-mapping analysis. All the fine-mapping results are reported in **Supplementary Table 6**.

#### Parameter optimization

FINEMAP and CAVIARBF use different default values for prior standard deviation of effect sizes (FINEMAP: 0.05; CAVIARBF: 0.1281429). We tuned different prior values ∈ {0.05, 0.12, 0.2, 0.3, 0.4, 0.5, 1, calculated from the data itself}, and observed different values severely affect the fine-mapping results for QTLs, however, no significant difference was observed for IMDs (**Supplementary Fig. 8**). Therefore, we set an acceptable prior for QTL to 0.3 and maintained the CAVIARBF default value (0.1281429) for IMD.

#### Definition of 95% credible set

To identify potential causal variants for each locus, we created 95% credible set for QTL and IMD separately, assuming a single causal variant per locus. We created 95% credible sets by ranking all variants in a locus based on the posterior probability, and including variants until the sum of posterior probabilities was ≥ 0.95.

#### Comparable IMD-QTL dataset for fine-mapping

We observed that a subset of IMD datasets had lower numbers of variants per genomic region compared to the BLUEPRINT data, especially in the case of the focused Immunochip content. In order to control for these variant density differences between IMD and QTL loci, and to ensure a fair comparison for fine-mapping, we considered only loci where at least 80% reciprocal overlap between the variants contained in each genomic interval. Further, given that fine-mapping methods are constrained by inherent power limitations of the data, in order to robustly compare credible sets between QTL and IMD, we considered only disease loci reaching genome-wide significant (*P* ≤ 5 × 10^−8^) levels of association in the available summary statistics. We further removed *GPR35* locus that are associated with AS, IBD, and UC and colocalised with mQTL, as it could not be fine-mapped using mQTL, although rs4676410 was the top variant in mQTL credible set with highest *PP*_*fm*_ (0.48).

### Comparison with IBD fine-mapping based on individual-level data

In a recent study, Huang and colleagues attempted to fine-map 94 IBD loci using high-density genotype data ^9^. Of these, 68 loci were found to contain a single independent association, while others contained multiple independent signals. We overlapped these loci with our QTL data and only considered the loci that meet the following criteria: (i) the disease loci showed strong colocalisation evidence (*PP*_*coloc*_ ≥ 0.98) with at least one of the QTLs in one cell type; (ii) selected QTL loci contained only one independent causal variant with MAF ≥ 5%; and (iii) both QTL and IBD fine-mapping credible set size ≤ 100 variants. Finally, we selected 32 loci full-filling above criteria. For each locus, we compared the reported credible set and the minimal credible set out of all QTLs (**Supplementary Fig. 9 and Supplementary Table 7**).

## Data availability

The BLUEPRINT phase 2 Genotype data (VCFs) have been deposited in the European Genome-phenome Archive (EGA) under accession EGAD00001005192. All the QTL summary statistics are available under EGAD00001005199 and EGAD00001005200.

## Acknowledgements

Kousik Kundu is supported by the NIHR CBR (Cardiovascular Theme). This study was conducted using the BLUEPRINT (http://www.blueprint-epigenome.eu/) data funded by EU FP7 High Impact Project BLUEPRINT (HEALTH-F5-2011-282510) and the Canadian Institutes of Health Research (CIHR EP1-120608). We thank Lu Chen and Valentina Iotchkova for the initial technical discussion on analysis strategy, and Katrina M de Lange for helping with IBD GWAS data. We sincerely thank Hilary Martin and Emma Davenport for their invaluable comments on the manuscript. We also thank Quan Lin for releasing the new BLUEPRINT phase 2 data through European Genome-phenome Archive (EGA), EMBL-EBI and acknowledge support from the Cambridge NIHR Biomedical Research Centre and the International Multiple Sclerosis Genetics Consortium (IMSGC). We also gratefully acknowledge Willem H. Ouwehand, Kate Downes as part of the National Health Service (NHS) Blood and Transplant for their contribution on volunteer recruitment and blood collections.

## Author contributions

K.K. and N.S. designed the study. K.K. and A.L.M. acquired the data, K.K performed the analysis. K.K., A.L.M., M.T., S.W., C.A.A., and N.S., interpreted the results. K.K., A.L.M., M.T., and N.S. wrote the manuscript. All authors read and approved the final version of the manuscript.

