## Supplementary Information for "Genetic associations at regulatory phenotypes improve fine-mapping of causal variants for twelve immune-mediated diseases"

**Address for correspondence:**

Prof. Nicole Soranzo
Human Genetics
Wellcome Sanger Institute
Genome Campus
Hinxton, CB10 1HH

*to whom correspondence should be addressed.

### Supplementary figures:

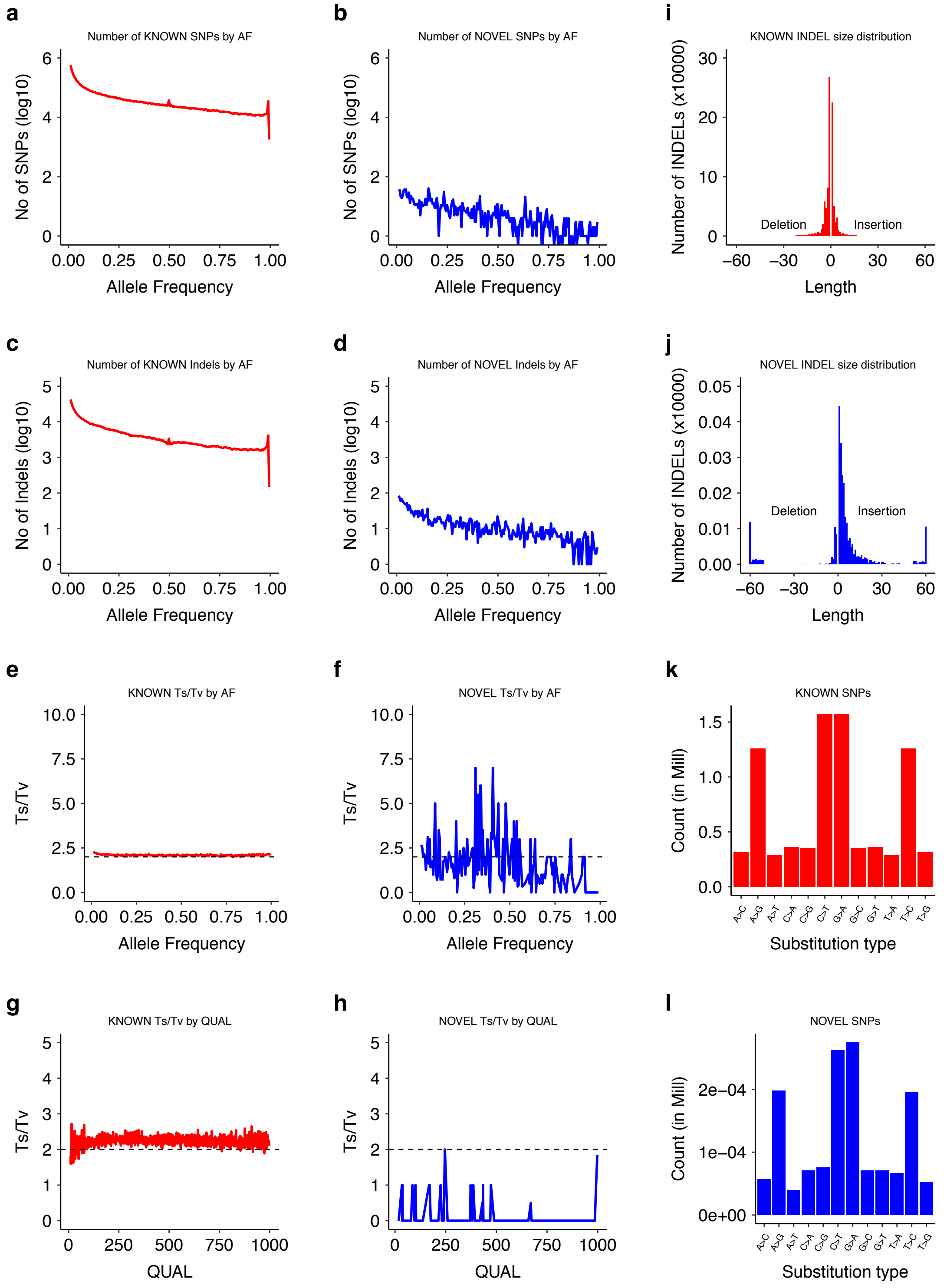


**Supplementary Fig. 1**: *Blueprint phase 2 variant statistics.* Variant statistics were generated using BCFTools (http://samtools.github.io/bcftools/). **a-d**, Total number of SNPs and INDELs by alternative allele frequency (AAF) split by known and novel (not present in dbSNP v149) sites. **e-h**, Transition/transversion ratio (Ts/Tv) by allele frequency (e,f) and by GATK quality score (g,h) split by known and novel. **i,j,** Size distribution shown separately for known and novel INDELs. **k,l,** Distribution of substitution types for known and novel SNPs.

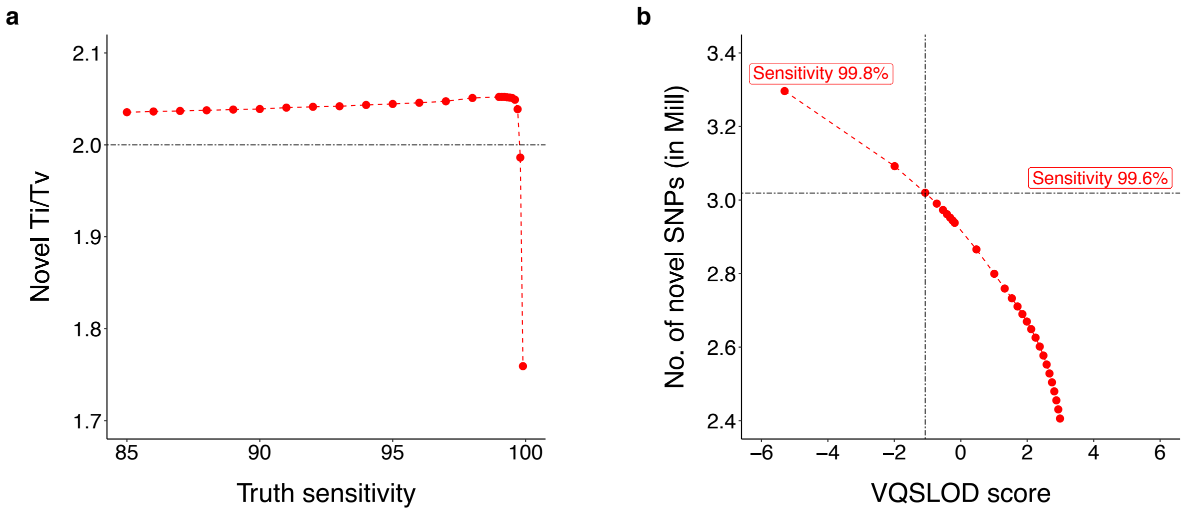


**Supplementary Fig. 2**: *GATK VQSR filter for variants.* **a**, We set the threshold at 2.0491 Transition/transversion (Ti/Tv) ratio for novel variants to achieve 99.6% truth sensitivity for SNPs. **b**, The corresponding VQSLOD score was -1.0707, which includes ∼3 million novel (not present in dbSNP v138) SNPs.

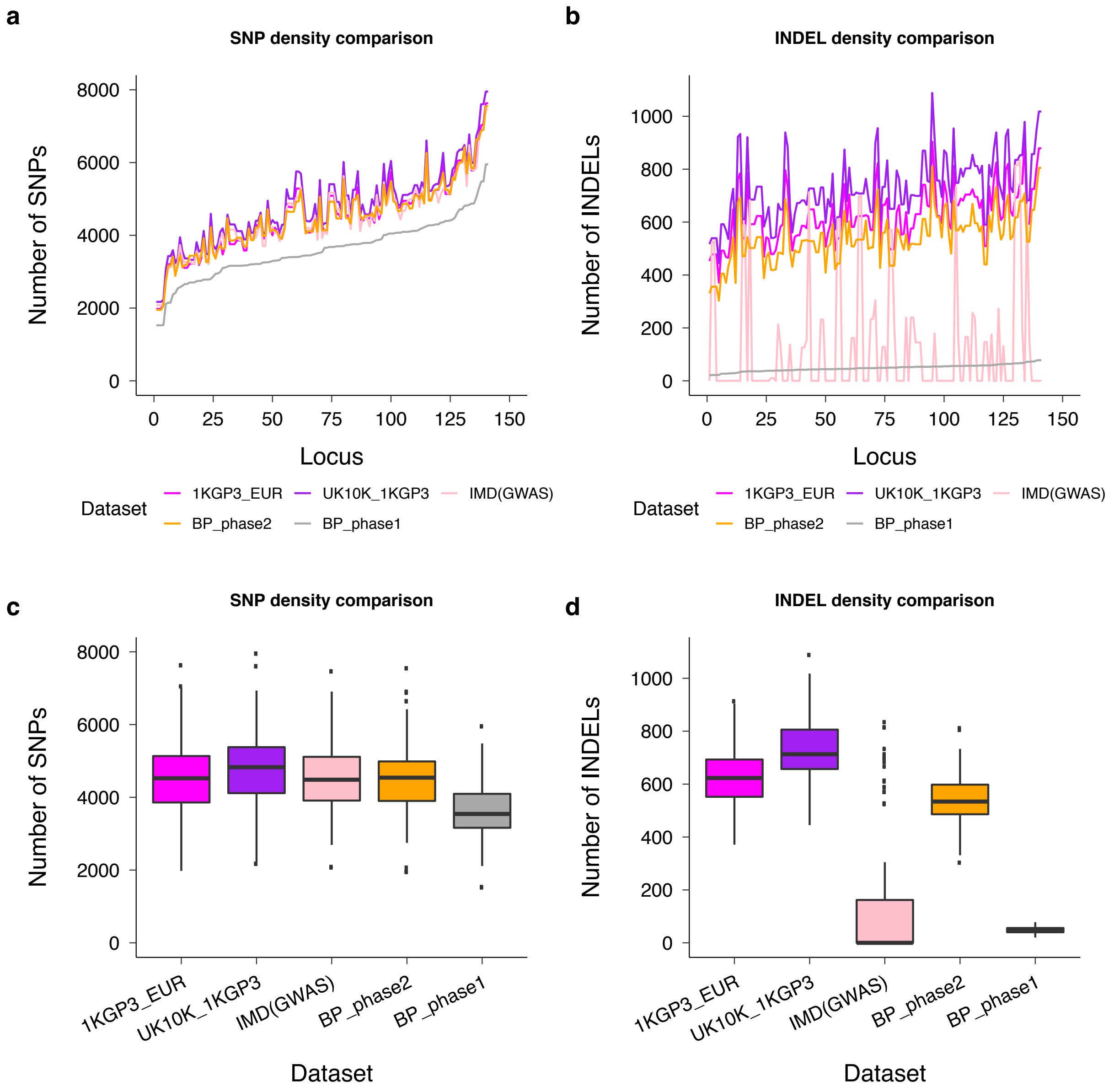


**Supplementary Fig. 3**: *Variant density comparison in the BLUEPRINT phase 1 and phase 2 datasets.* **a,b**, Total number of SNPs (a) and INDELs (b) in different datasets and for each IMD locus investigated. For a fair comparison, we only considered IMD loci that have at least 80% reciprocal overlap with QTL loci. The figures indicate that the phase 2 data includes larger numbers of variants compared to the phase 1 data, comparable with other whole-genome sequencing datasets. In phase 2, importantly, a significant improvement was achieved for INDEL density (b) than for SNP density (a). **c,d**, Distribution of number of variants in different datasets. Overall, phase 2 data captured more INDELs compared to phase 1 and IMD data.

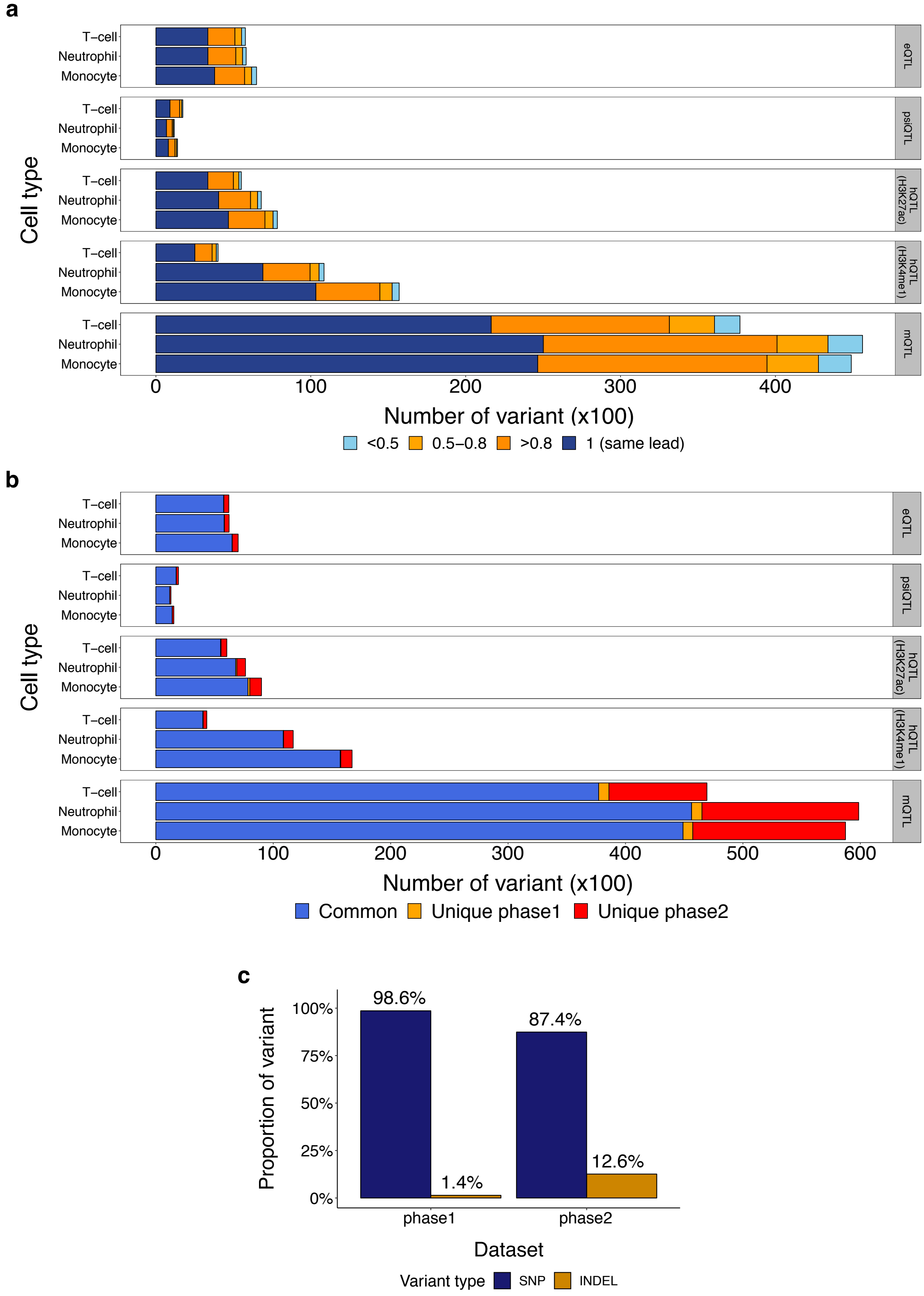


**Supplementary Fig. 4**: *Quantitative trait locus (QTL) comparison for BLUEPRINT phase 1 and phase 2 data.* **a**, Most of the lead variants are either the same or highly correlated ($r^2 > 0.8$) in both datasets. **b**, We obtained comparable QTL discovery (5% gFDR) in phase 1 and phase 2 datasets; the slightly higher number of QTLs achieved in phase 2 is due to a less stringent multiple test correction (Eigen p-value was used to calculate gFDR in phase 2 data, where Bonferroni p-value was used for phase 1 data). **c**, Proportion of SNP and INDEL lead variants in phase 1 and phase 2 data. As we captured more INDELs in our call set, the proportion of INDELs in phase 2 data is markedly higher.

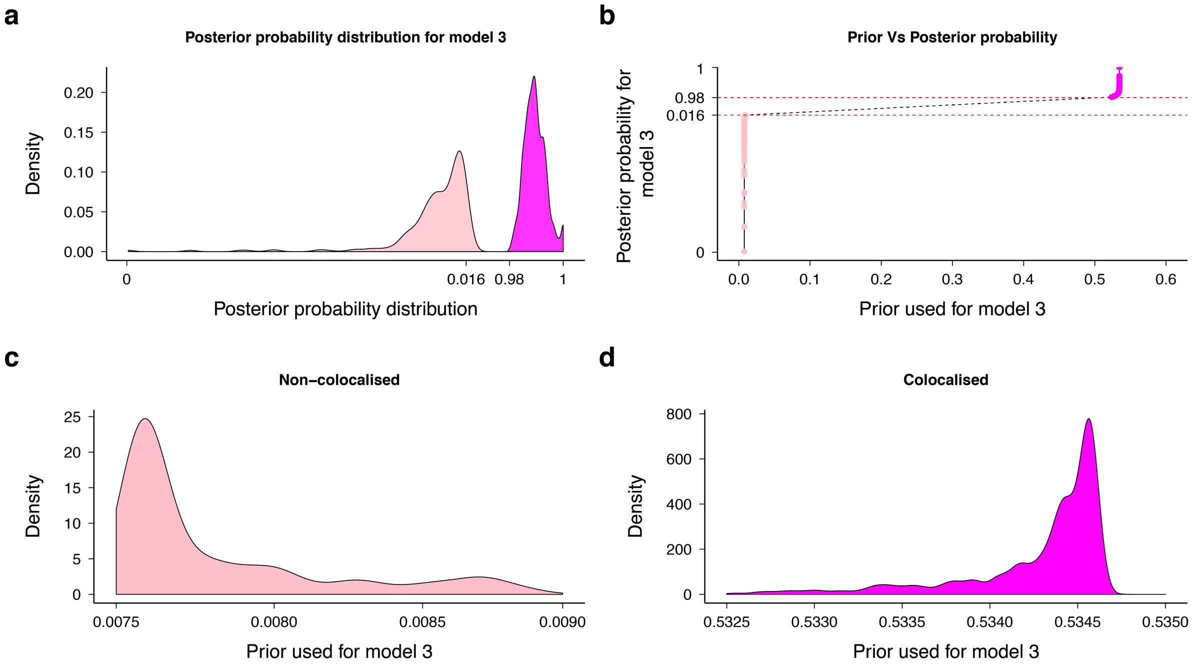


**Supplementary Fig. 5**: *Colocalisation prior distribution.* We used gwas-pw for colocalisation test[1]. **a,** Posterior probability distribution for model 3, supporting colocalisation. Two very distinct distributions were observed for the loci that colocalised (magenta; model 3) and the loci that do not colocalised (pink; strongly supporting model 4 as linkage evidence). **b,** Scatter plot between priors used and posterior probabilities for model 3. **c,d,** Prior distributions for model 3 plotted for colocalisaed and non-colocalised loci.

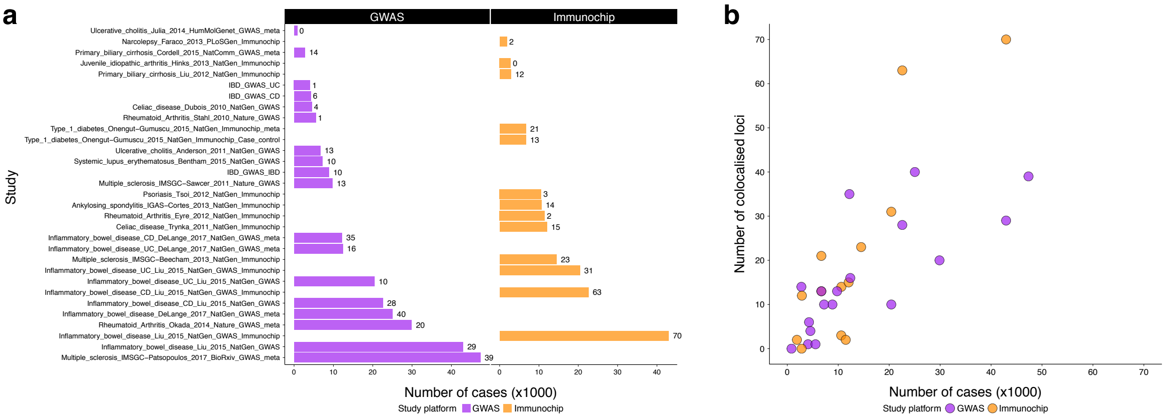


**Supplementary Fig. 6**: *Number of colocalised loci per study.* **a,** Sample size (cases) per study that have been used for colocalisation. Numbers next to each bar represent the number of colocalised loci ($PP_{coloc} \geq 0.98$) in respective studies. **b,** Correlation plot between sample size and number of colocalised loci. Different datasets (i.e., GWAS and immunochip) are depicted in different colors.

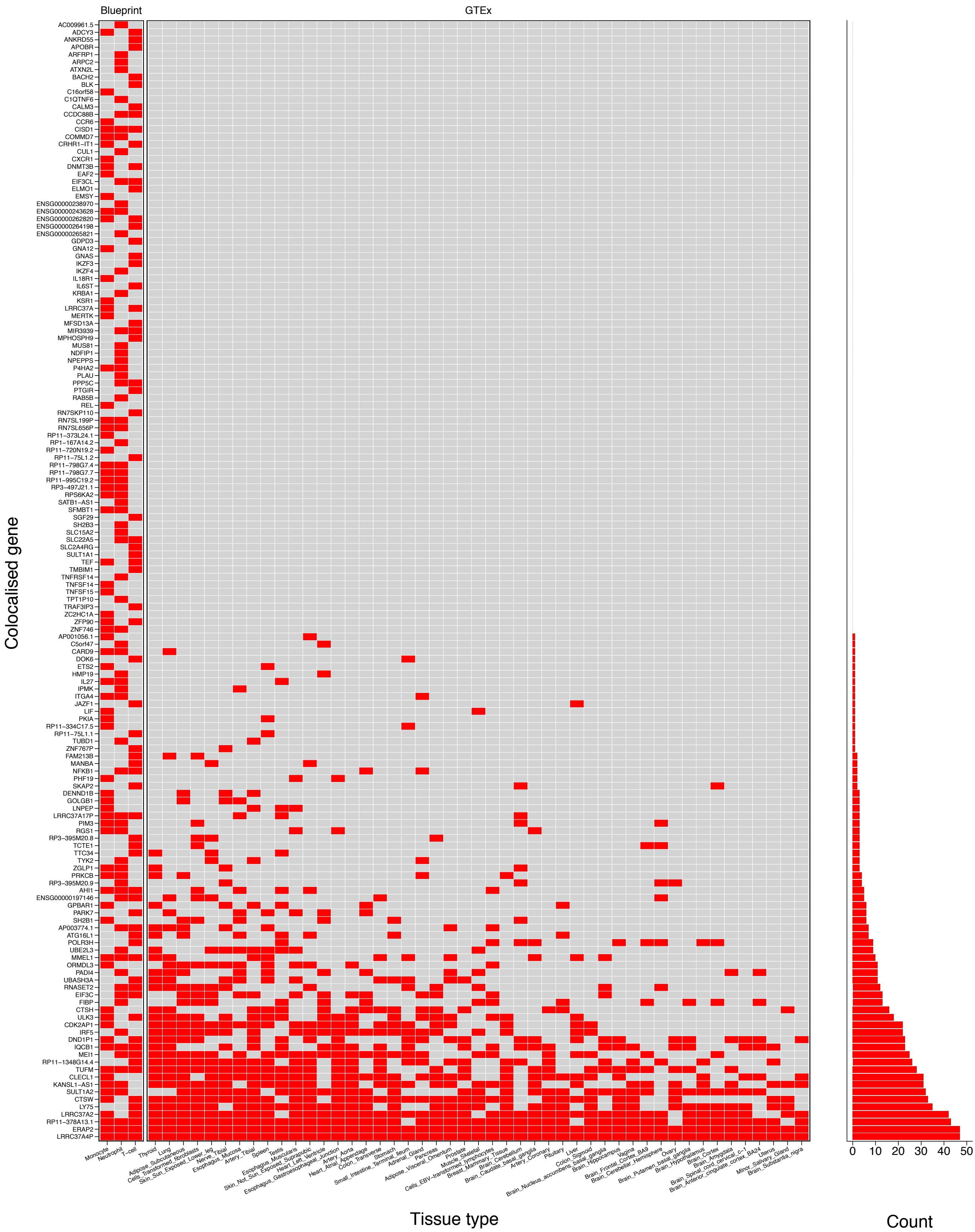


**Supplementary Fig. 7**: *Expression QTLs (eQTLs) overlapping with 47 GTEx (v7) tissues.* We searched sentinel BLUEPRINT leads and their proxies ($r^2 \geq 0.8$) against GTEx eQTLs (excluded "Whole Blood"). We assessed immune cell type specificity of 150 genes that colocalised with IMD signals. Left panel: heatmap describing gene-by-gene overlap between the two eQTL datasets (red: overlap; grey: non-overlap). The majority of eQTLs (82 genes) are specific to one or more of the three immune cell types, a smaller number of eQTLs (68 genes) map to multiple tissues. Right panel: number of GTEx tissues that share an eQTL signal with BLUEPRINT data.

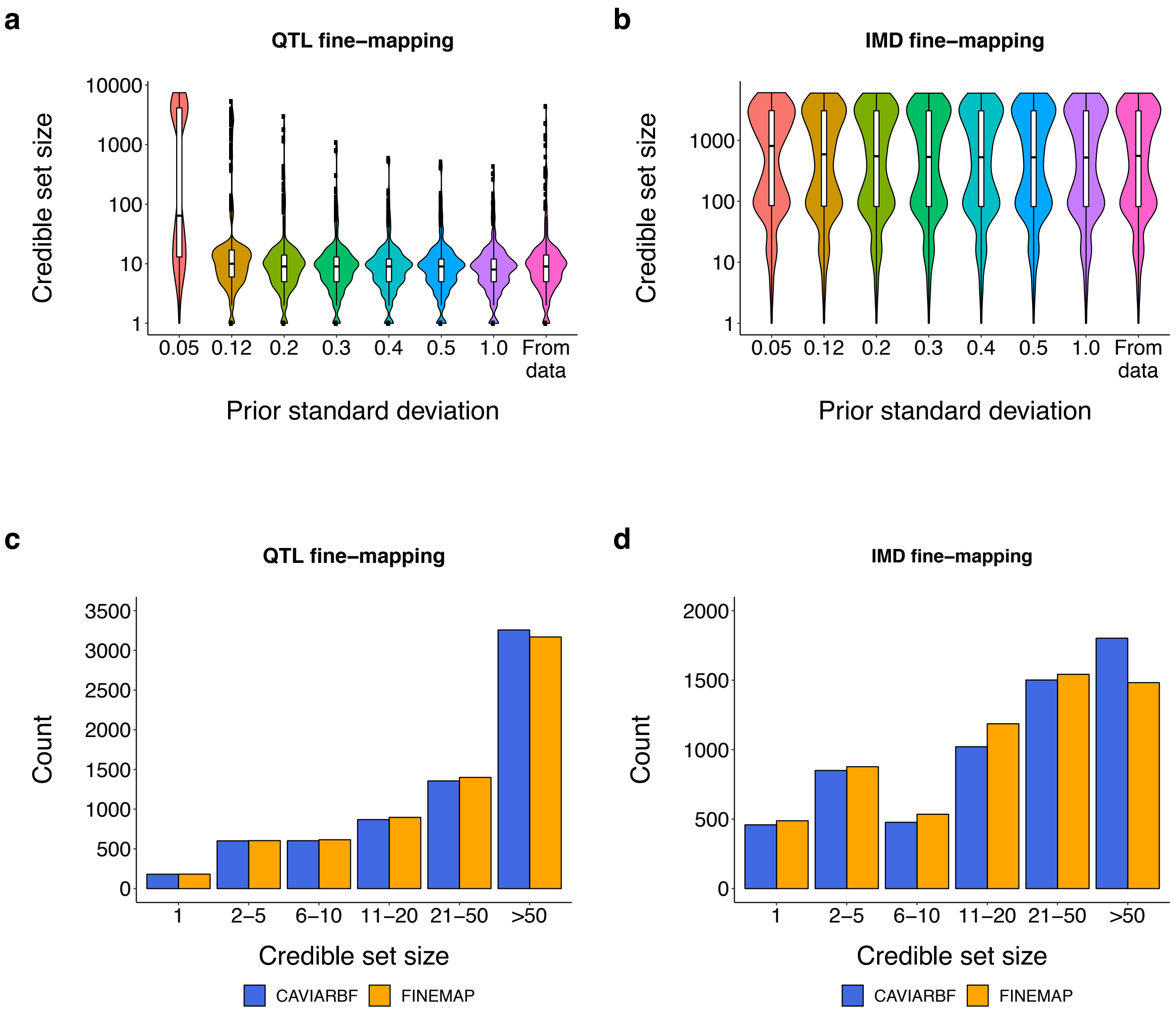


**Supplementary Fig. 8**: *Parameter optimization and comparison of different fine-mapping methods.* **a,b**, Distribution of credible set sizes in the QTL and IMD fine-mapping experiments, plotted as a function of prior standard deviation of effect sizes (default values for FINEMAP and CAVIARBF are 0.05 and 0.1281429, respectively). We observed the fine-mapping credible sets decrease for QTL loci, while IMD credible set size distribution does not decrease with increasing priors. We set prior standard deviation for QTL and IMD loci at 0.3 and 0.1281429 (default value of CAVIARBF), respectively. **c,d**, Comparison of FINEMAP and CAVIARBF credible set size in fine-mapping experiments. Credible sets were binned based on the number of variants. No significant difference was observed when comparing the FINEMAP and CAVIARBF results on both QTL and IMD loci.

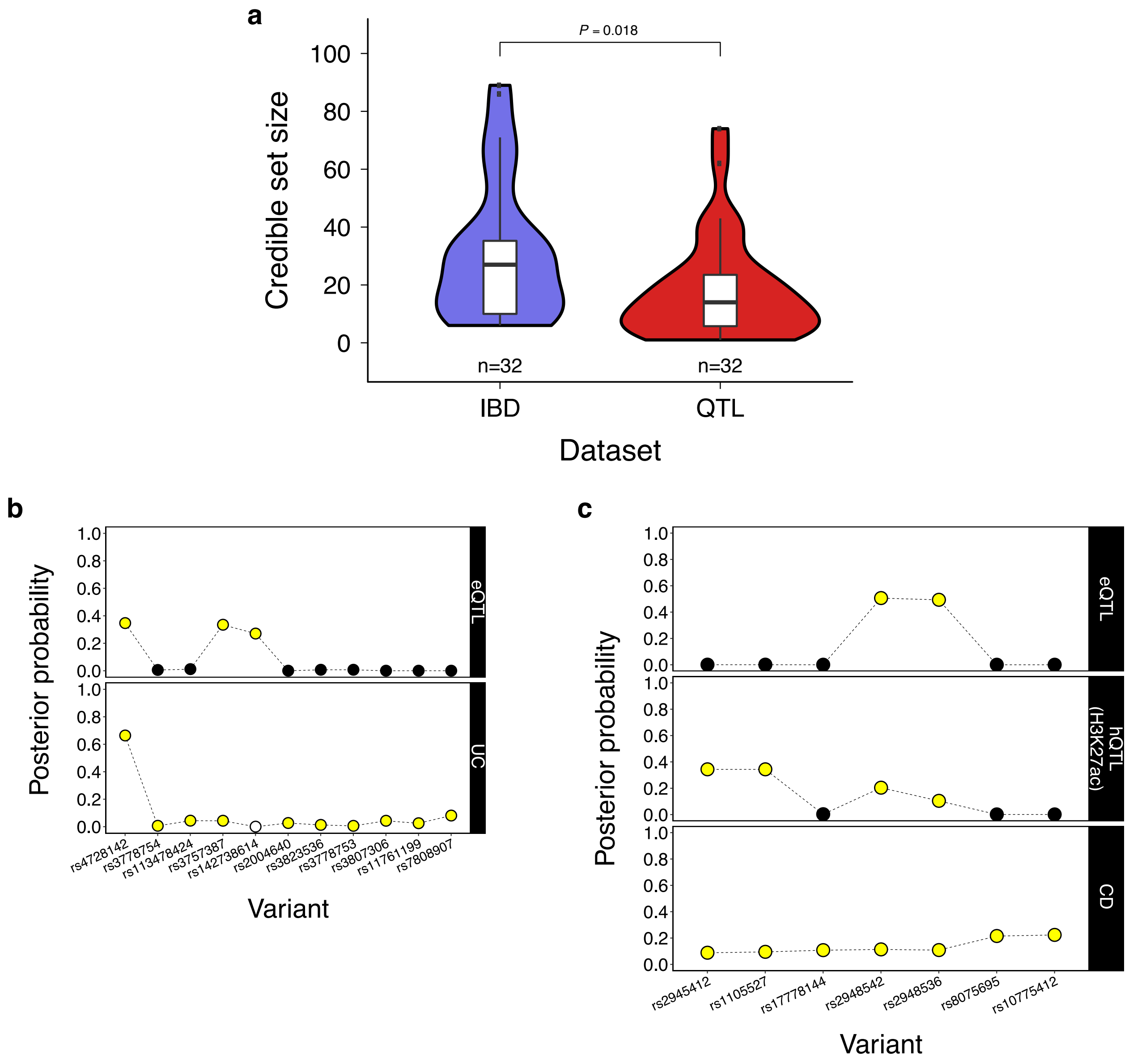


**Supplementary Fig. 9**: *Comparison with publicly available IBD fine-mapping data.* **a,** Credible set comparison between QTLs (minimal credible set out of five regulatory traits and three cell types) and recently published IBD fine-mapping data[2] for 32 colocalised loci that meet our criteria (**Methods**). The figure indicates that on average QTLs achieve smaller credible set (mean = 18.1) compared to IMD (mean = 30.81; t-test $P = 0.018$). **b,c,** Posterior probability comparison for two exemplar IBD loci: *IRF5* (b) and *KSR1* (c). Credible set variants are in yellow, while variants that were included in the fine-mapping analysis but which are not part of the credible set are in black. Variant rs142738614 (white) was not part of the IBD credible set; we could not check whetehr the variant was tested for fine-mapping analysis in Huang et a.[2] due to lack of data availability. In both cases, QTLs yielded smaller credible set compared to IMDs. For *IRF5*(a), the eQTL fine-mapping credible set contains three variants, while UC credible set contains ten[2]. rs4728142 achieved the highest posterior probability in both datasets. Interestingly, one of the three variants in the eQTL credible set (INDEL: rs142738614) was not part of the disease credible set. For *KSR1*, fine-mapping based on CD yielded a seven variant credible set[2], compared to a credible set of four variants for hQTL (H3K27ac) and two for eQTL. All QTL credible set variants are included in the disease fine-mapping credible set.

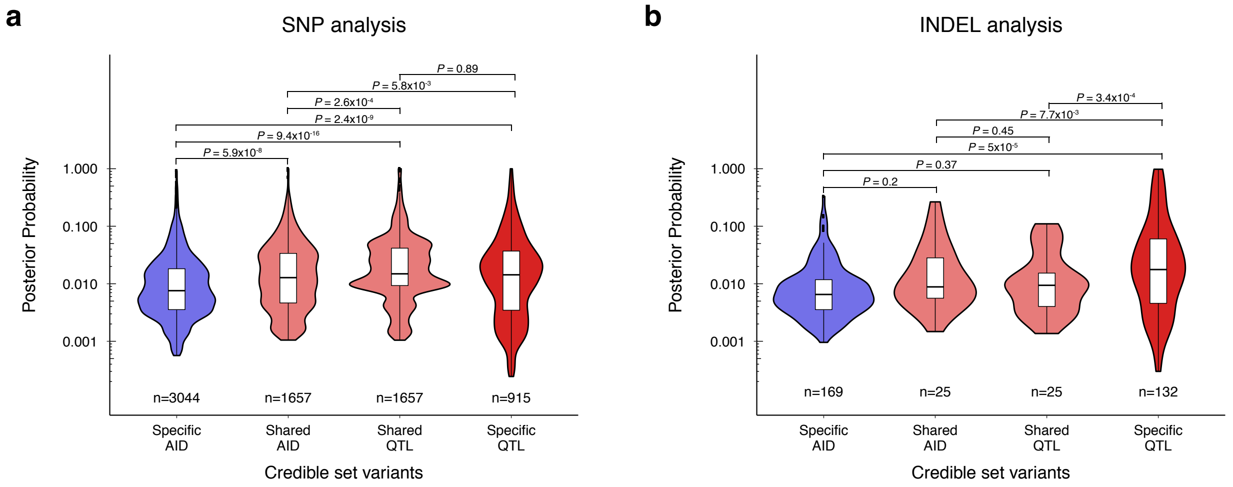


**Supplementary Fig. 10**: *Distribution of posterior probabilities for SNPs and INDELs that were shared (intersection) and specific to QTL and IMD loci.* **a,b**, Violin plots illustrate the posterior probability ($PP_{fm}$) distributions for SNPs (a) and INDELs (b) discovered by both the IMD and QTL analysis (intersection set), or uniquely contained in the IMD or QTL credible sets (IMD-specific and QTL-specific). A t-test was used to calculate significance values ($P$).

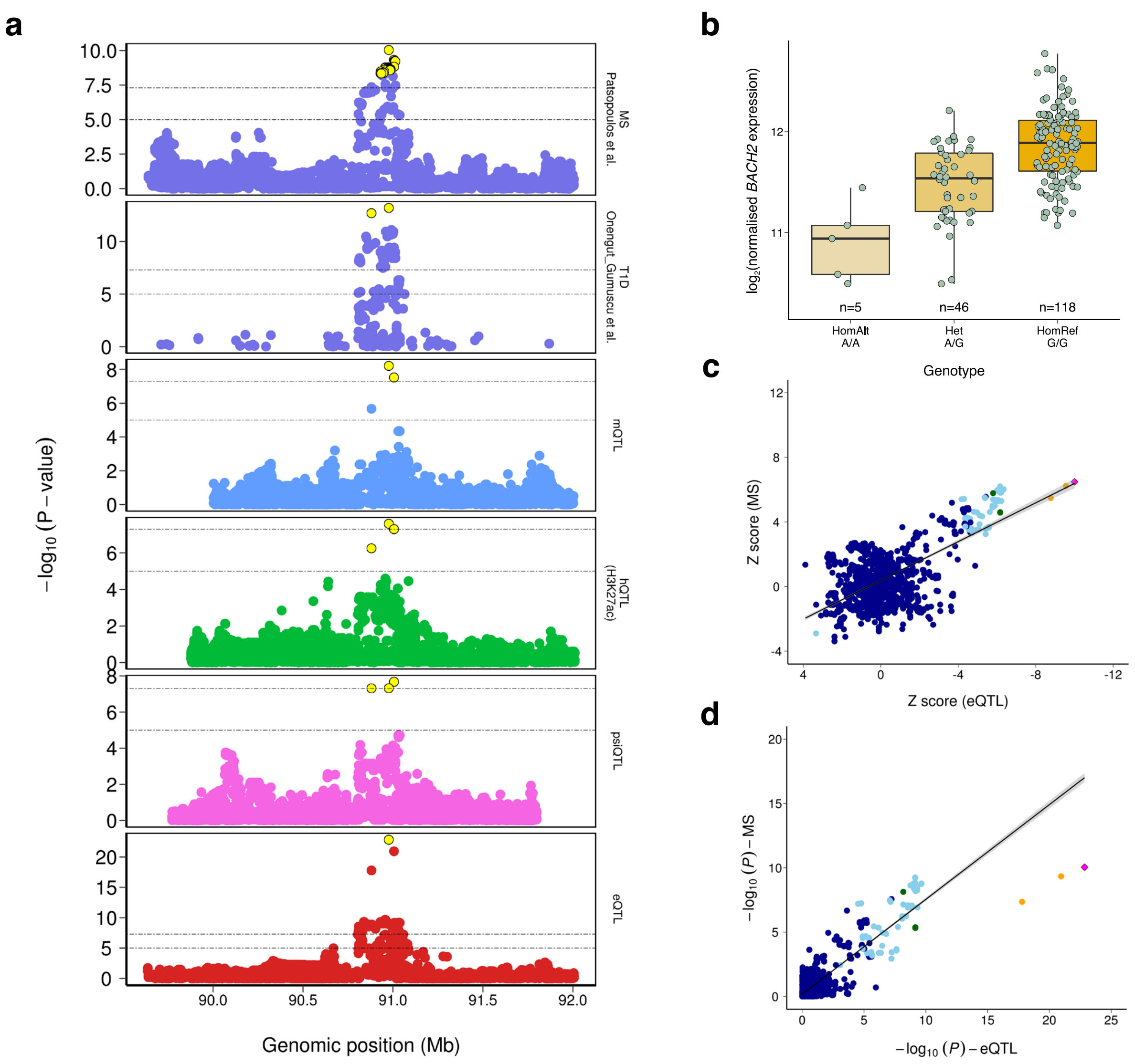


**Supplementary Fig. 11**: *Fine-mapping of the BACH2 locus.* The risk allele A of the intronic variant rs72928038, which is known to be associated with MS[3] and T1D[4], and lies in the intron closest to the Transcriptional Start Site (TSS) of *BACH2*. **a,** A colocalisation plot using IMD and QTL data with 250kb flanking region surrounding the sentinel SNP (rs72928038). This plot illustrates the locus was strongly colocalised ($PP_{coloc} \geq 0.98$) between IMDs and regulatory QTLs (i.e., eQTL, psiQTL, hQTL (H3K27ac) and mQTL) in CD4+ T-cells. Yellow circles represent the variants present in the respective 95% credible sets. **b,** The figure indicates the effect allele (A) of the sentinel variant (rs72928038) decreasing *BACH2* expression level. **c,d,** Z-score and p-value correlation plot between MS and eQTL. The figures indicates the colocalisation was effective, showing both are the same signal and the variants effects (rs72928038:A) are in the opposite directions. The case of the *BACH2* gene is a paradigmatic example. This gene is a key regulator of the final stages of B-cell and T-cell differentiation and prevents inflammatory disease by controlling the balance between tolerance and immunity[5,6]. Furthermore, a human immunodeficiency and autoimmunity syndrome (BRIDA) has been described in cases of *BACH2* haploinsufficiency[7]. We identified that the risk allele was associated with diminished *BACH2* expression (b) in addition to reduced enhancer binding (colocalised with H3K27ac QTL) and increased methylation signal. The epigenetic changes and the subsequent effect on *BACH2* gene expression point to the presence of a transcription factor binding in the area of the SNP. In line with this hypothesis, experiments carried out in mice CD4+ T cells have demonstrated the binding of Menin, a transcription factor that subsequently recruits Histone-Acetyltransferases (HATs), to the intron close to the *BACH2* TSS[8].

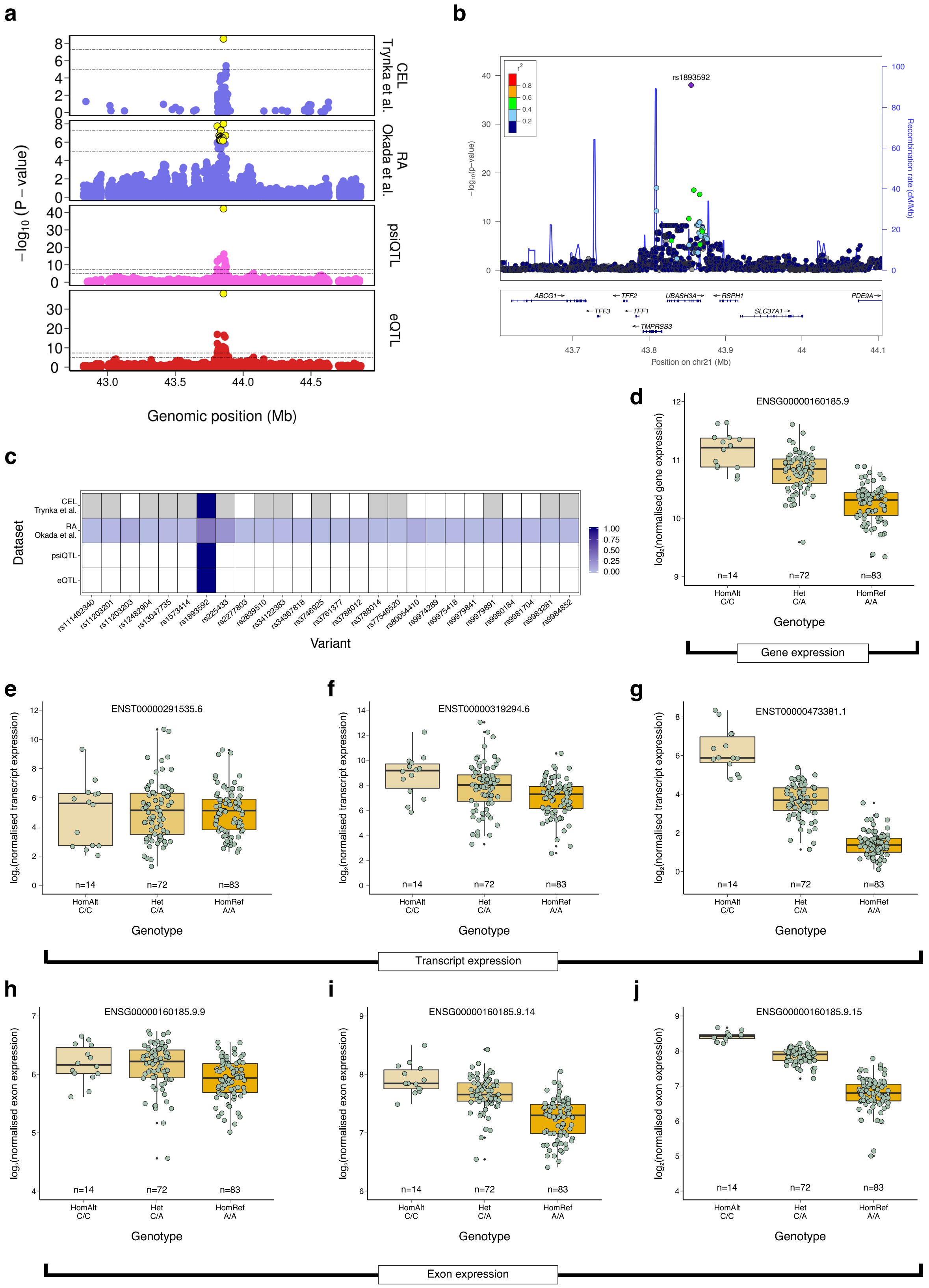


**Supplementary Fig. 12**: *Fine-mapping of the UBASH3A locus.* The rs1893592 is a *UBASH3A* intronic SNP and a risk allele for CEL[9], RA[10] and Primary sclerosing cholangitis (PSC; not included in our analysis)[11]. This variant was also reported as an eQTL and splicing QTL[12,13] in blood cells. In our data, rs1893592 was also seen to be an eQTL and psiQTL in CD4+ T-cells. **a,** Colocalisation

**Supplementary Fig. 12**: (continued..) plot indicates the locus is confidently colocalised ($PP_{coloc} \geq 0.98$) between IMDs and QTLs. Yellow circles represent the variants present in the respective credible sets. **b,** A locuszoom plot using eQTL data with 250kb flanking region surrounding the sentinel SNP (rs1893592). **c,** Heatmap of posterior probability ($PP_{fm}$) of the variants in the respective credible sets (colour intensity: $PP_{fm\_smallest}$ - light blue to $PP_{fm\_largest}$ - deep blue). White colour indicates the variants are not part of the respective credible sets, while grey colour indicates that the variants were not present in the respective summary statistics. The lead variant (rs1893592) appeared to be the single fine-mapped variant ($PP_{fm} \sim 1$) for eQTL, psiQTL and CEL. The locus could not be fine-mapped confidently with RA summary statistics, yielded 26 variants in the credible set, although rs1893592 achieved highest posterior probability ($PP_{fm} = 0.37$). As the variant density was very low in CEL summary statistics many variants were not included in disease summary statistics and hence those variants could not be tested for fine-mapping (grey). The figure illustrates the power of QTLs to resolve causal variants compared to disease summary statistics. **d-j,** *UBASH3A* expression levels (d), transcript expression levels (e-g), and exon expression levels (h-j) in CD4+ T-cells by rs1893592 genotypes. *UBASH3A* gene, transcripts and exons quantification are given as a function of genotype at the sentinel variant rs1893592. Our analysis supports previous observations that rs1893592 results in a splicing effect that ultimately affects overall *UBASH3A* expression by increasing the level of a transcript that does not produce the protein[14].

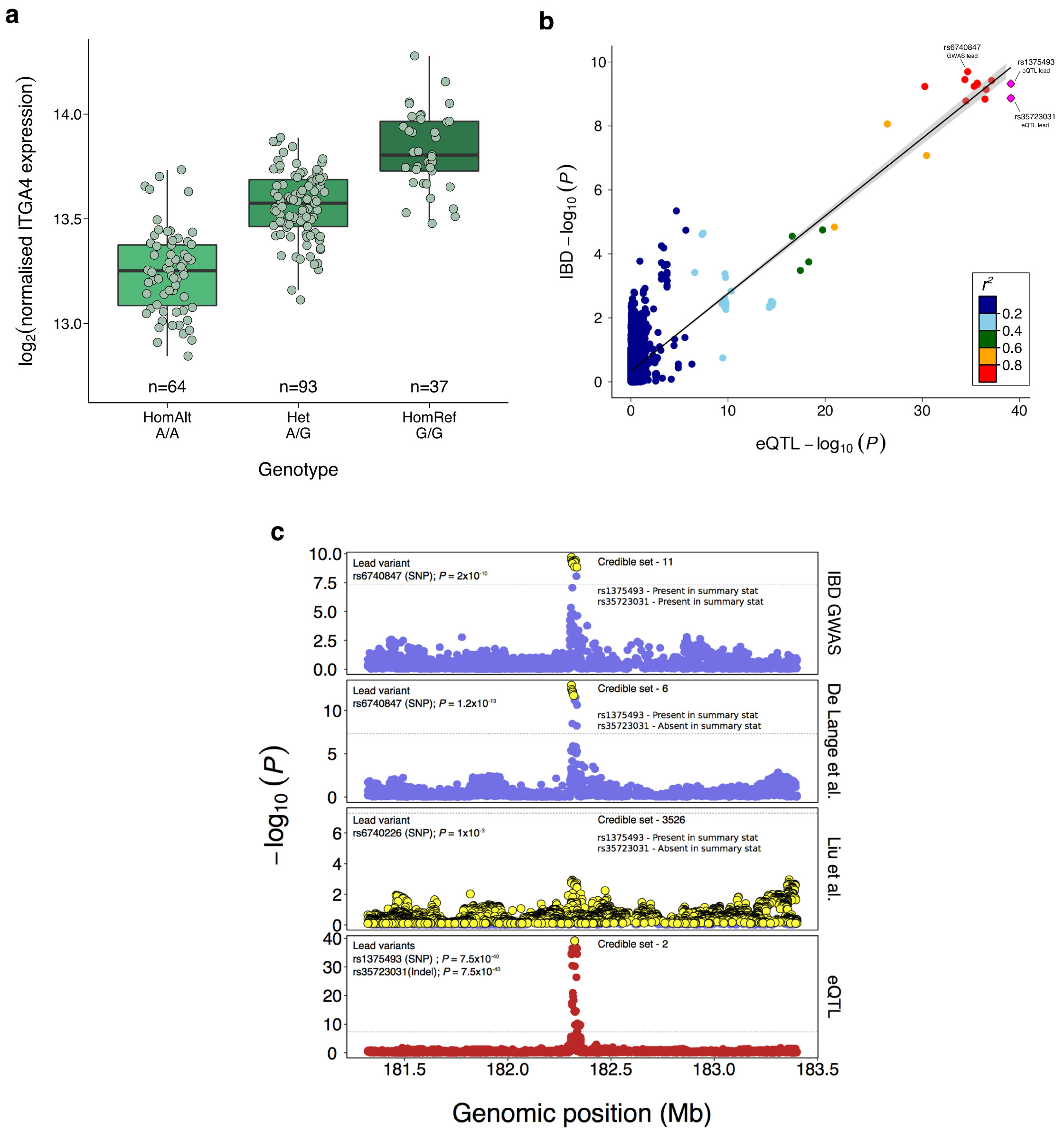


**Supplementary Fig. 13**: *ITGA4 locus analysis.* **a**, *ITGA4* expression levels in monocytes by rs1375493 genotypes. **b**, Correlation of IMD and eQTL p-values for all variants in a 1Mb region surrounding the sentinel SNP. **c**, Fine-mapping results for the *ITGA4* locus using different GWAS summary statistics[15,16]. The 95% credible set variants are denoted as yellow circles. Note that the locus could not be fine-mapped using the summary statistics published by Liu et al.[16] due to low power.

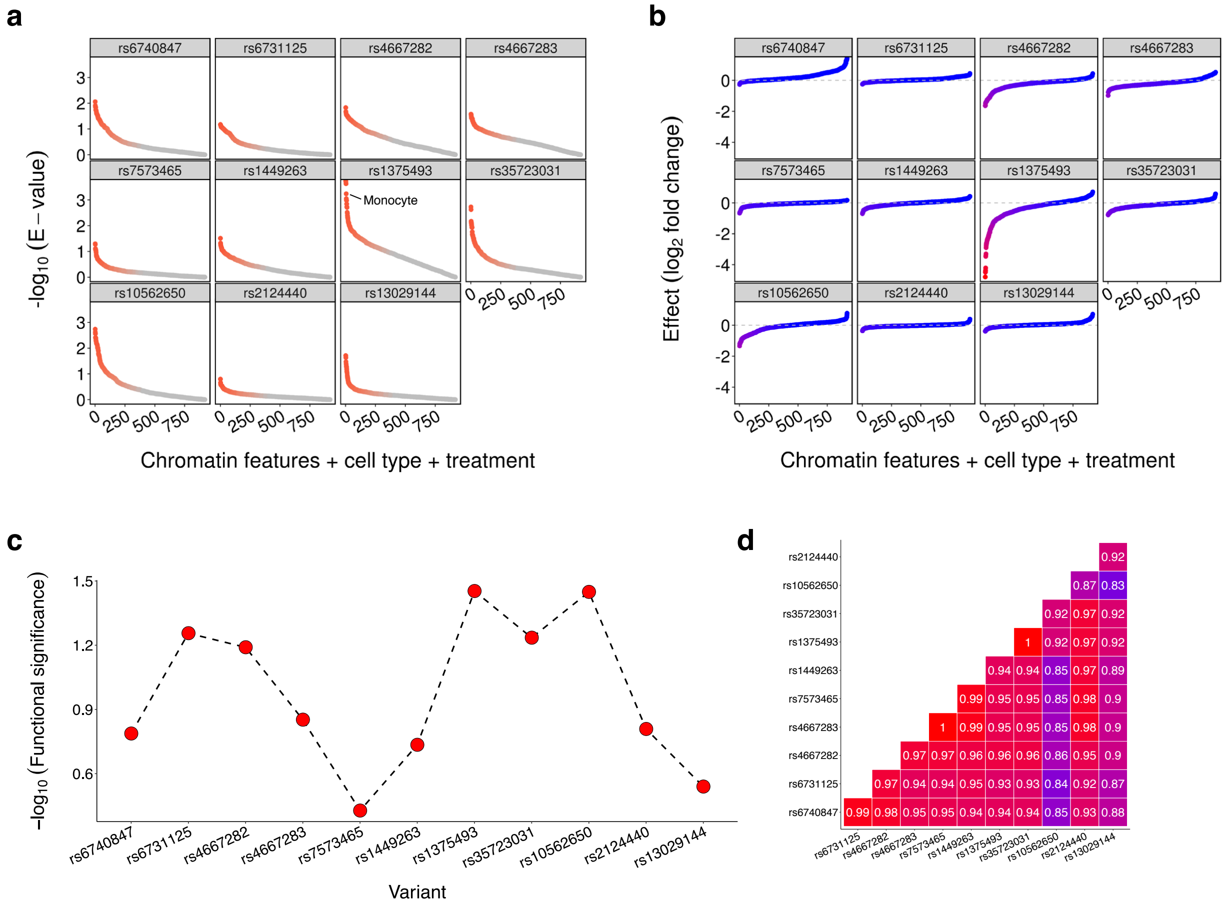


**Supplementary Fig. 14**: *Prediction of chromatin effects at the ITGA4 locus.* We predicted the chromatin effects of 11 variants at the *ITGA4* locus using DeepSEA [17], which is a deep-learning based prediction method. DeepSEA predicts a total 919 cell type and condition specific chromatin features (chromatic feature name + cell type + treatment) for each variant of interest. **a**, Chromatin feature probability log fold changes for each of the 11 variants. **b**, E-value for the effect of the chromatin features. **c**, Functional significant score for each variant. The figures show our lead variant rs1375493 was predicted to be more functionally significant in all the matrices compared to other variants, specifically in monocytes. **d**, Pairwise linkage disequilibrium (LD) of all 11 variants (credible set of IBD GWAS data) calculated using BLUEPRINT data.

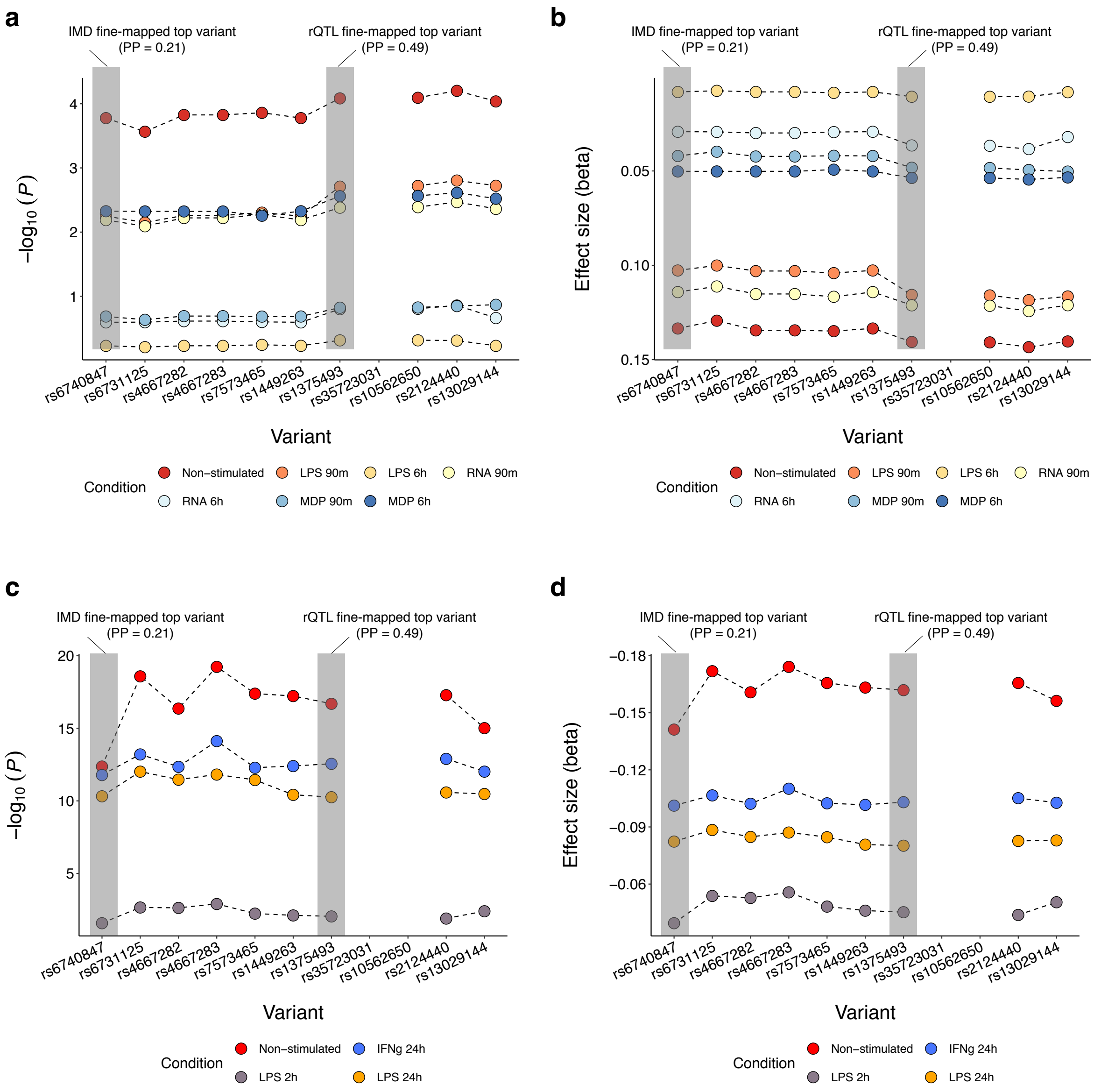


**Supplementary Fig. 15**: *Associations of the ITGA4 locus in stimulated monocytes.* **a-d**, Association p-values of 11 credible set variants in different stimulated monocytes assayed in Kim-Hellmuth et al.[18] (a,b) and Fairfax et al.[19] (c,d) . In all conditions, our most-associated SNP rs1375493 showed greater significance than the IBD sentinel SNP rs6740847. Note that out of two INDELs in the credible set, rs35723031 (achieved highest $PP_{fm}(0.49)$ along with rs1375493 in eQTL data) was not tested in both datasets, whereas other INDEL: rs10562650 tested in Kim-Hellmuth et al.[18] only.

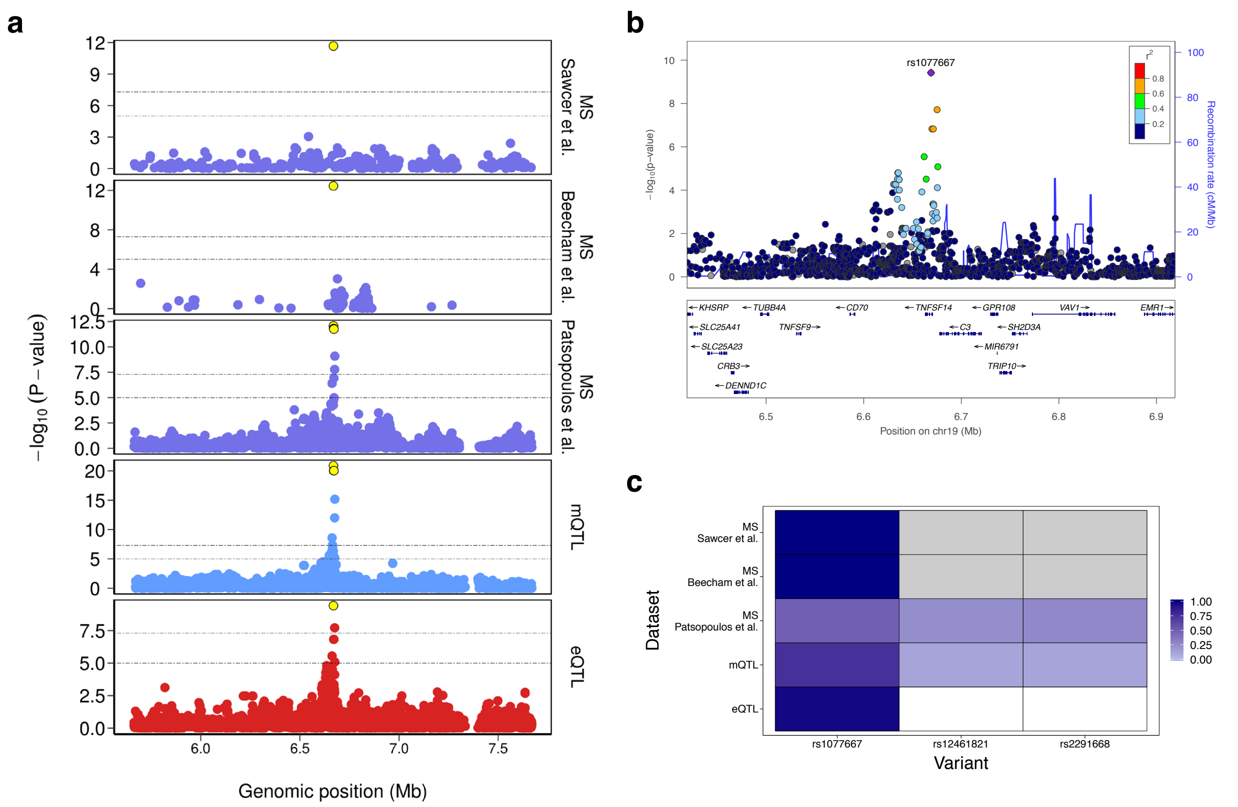


**Supplementary Fig. 16**: *Fine-mapping of the TNFSF14 locus.* The intronic variant rs1077667(T), is a protective allele for MS[3,20,21]. In our data, the variant is an eQTL, increasing the expression of *TNFSF14* and mQTL that reduces the signal for an overlapping methylation probe in monocytes. Importantly, this is observed in myeloid cells but not CD4+ T cells. **a,** The locus was highly colocalised ($PP_{coloc} \geq 0.98$) between MS and QTLs. **b,** A locuszoom plot using eQTL data with 250kb flanking region surrounding the sentinel SNP (rs1077667). **c,** Heatmap of posterior probability ($PP_{fm}$) of the variants in the respective credible sets (colour intensity: $PP_{fm\_smallest}$ - light blue to $PP_{fm\_largest}$ - deep blue). White colour indicates the variants are not part of the respective credible sets, while grey colour indicates that the variants were not present in the respective summary statistics. The lead variant (rs1077667) appeared to be the single fine-mapped variant ($PP_{fm} = 0.97$) for eQTL. The variant also achieved highest posterior probability in all other datasets, indicating most likely causal variant. As the variant density was very low in two MS summary statistics[20,21], many variants were not included in respective studies and hence rs12461821 and rs2291668 (part of the credible sets for mQTL and MS[3]) could not be tested for fine-mapping (grey). *TNFSF14* emerges as a particularly interesting gene, highlighted by our colocalisation analysis, as it modulates the HVEM-BTLA immune checkpoint[22].

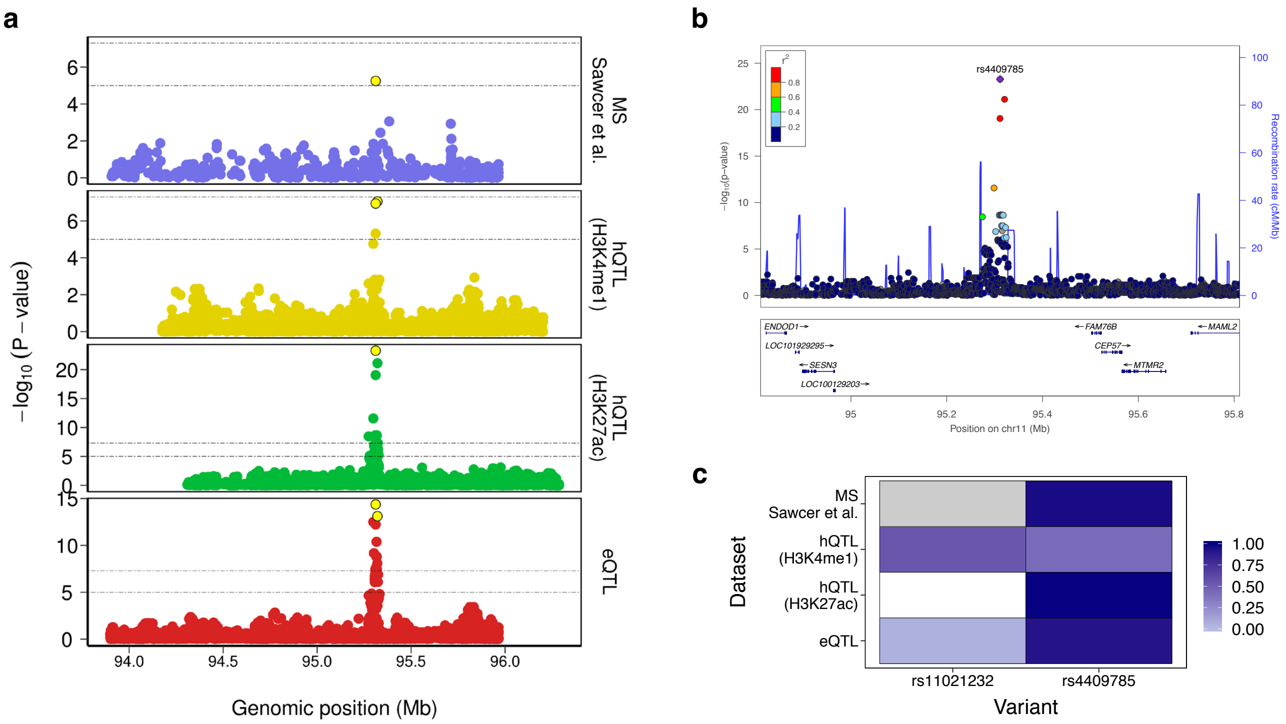


**Supplementary Fig. 17**: *Fine-mapping of the SESN3 locus.* **a,** The MS[21] risk allele, rs4409785(C), colocalised with an eQTL for *SESN3*, an antioxidant gene involved in the clearance of reactive oxygen species derived from RAS signalling. *SESN3* expression decrease was associated with the C-allele of rs4409785 and a decrease of H3K27ac signal. Note that the variant was not genome-wide significant in available summary statistics ($P = 5.626 \times 10^{-06}$)[21]. **b,** A locuszoom plot using eQTL data with 500kb flanking region surrounding the sentinel SNP (rs4409785). **c,** Heatmap of posterior probability ($PP_{fm}$) of the variants in the respective credible sets (colour intensity: $PP_{fm\_smallest}$ - light blue to $PP_{fm\_largest}$ - deep blue). White colour indicates the variants are not part of the respective credible sets, while grey colour indicates that the variants were not present in the respective summary statistics. This signal was specific to CD4+ T-cells in our analysis and we did not observe any sharing with other GTEx brain tissue-specific eQTLs, which argues against a possible role of the SNP in neurons exacerbating the toxic effects of reactive oxygen species in the context of Multiple Sclerosis.

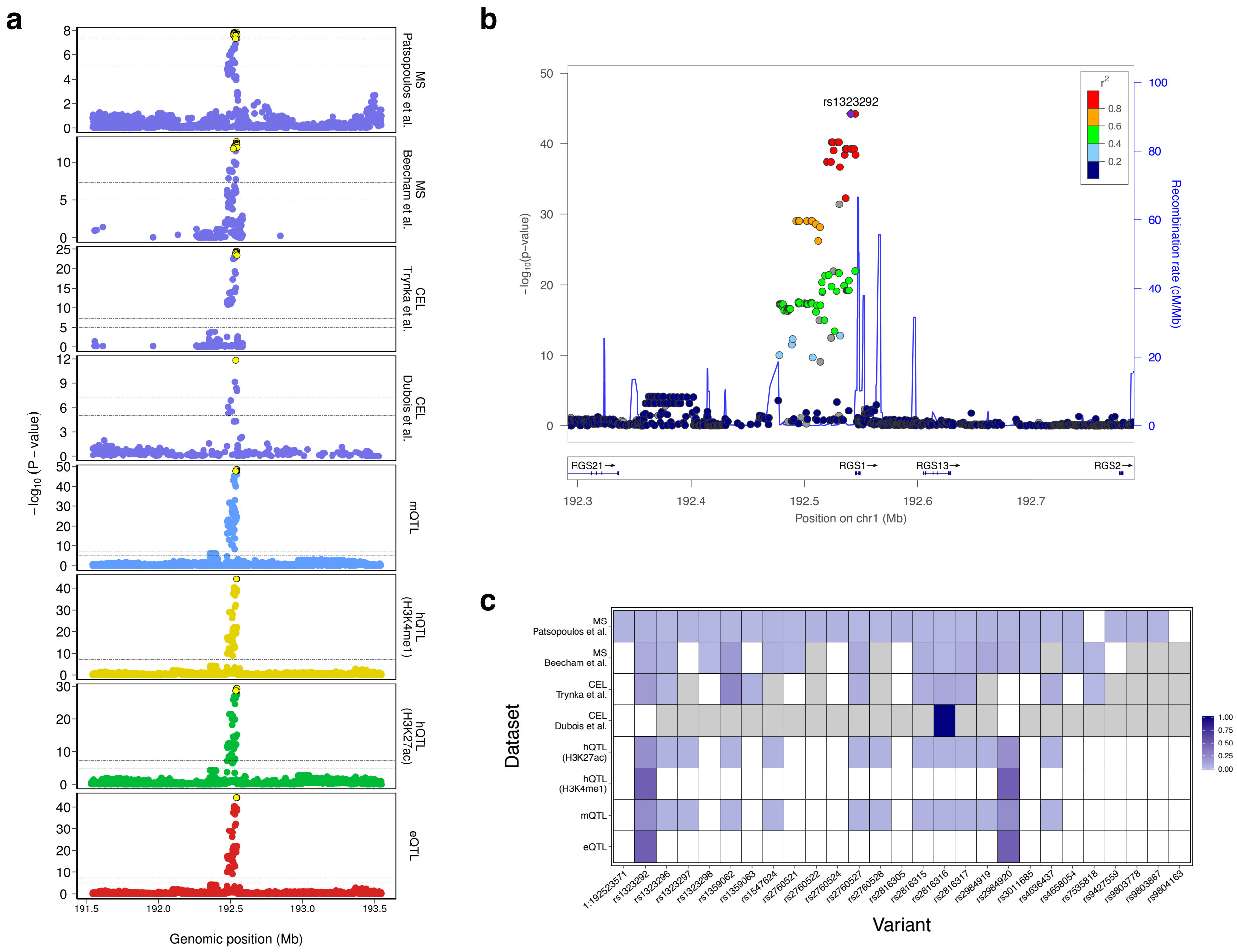


**Supplementary Fig. 18**: *Fine-mapping of the RGS1 locus. RGS1* is a GAP factor expressed by all leukocytes and has been implicated in the entry of B Lymphocytes into Lymph Nodes[23]. The locus is associated with both MS[3] and CEL[9,24], and colocalised with eQTL (for *RGS1*), hQTLs (H3K4me1 and H3K27ac), and mQTL in monocytes and neutrophils. **a**, Colocalisation plot of the locus, depicting regional association for disease locus (blue), eQTL (red), mQTL (lightblue), hQTL (H3K27ac in green and H3K4me1 in yellow) in neutrophils. **b**, A locuszoom plot for the *RGS1* locus using eQTL (neutrophil) data. **c**, Heatmap of posterior probability of variants derived from different credible sets. White boxes indicate the variants are not part of the respective credible sets and grey boxes indicate the variants are actually not included in the respective disease summary statistics. From QTL fine-mapping, rs1323292 and rs2984920 ($r^2$ = 1 in BLUEPRINT data; both have $PP_{fm}$ = 0.5) appear to be most probable causal variants among others. These variants are also part of the IMD credible sets for MS[3] and CEL[9].

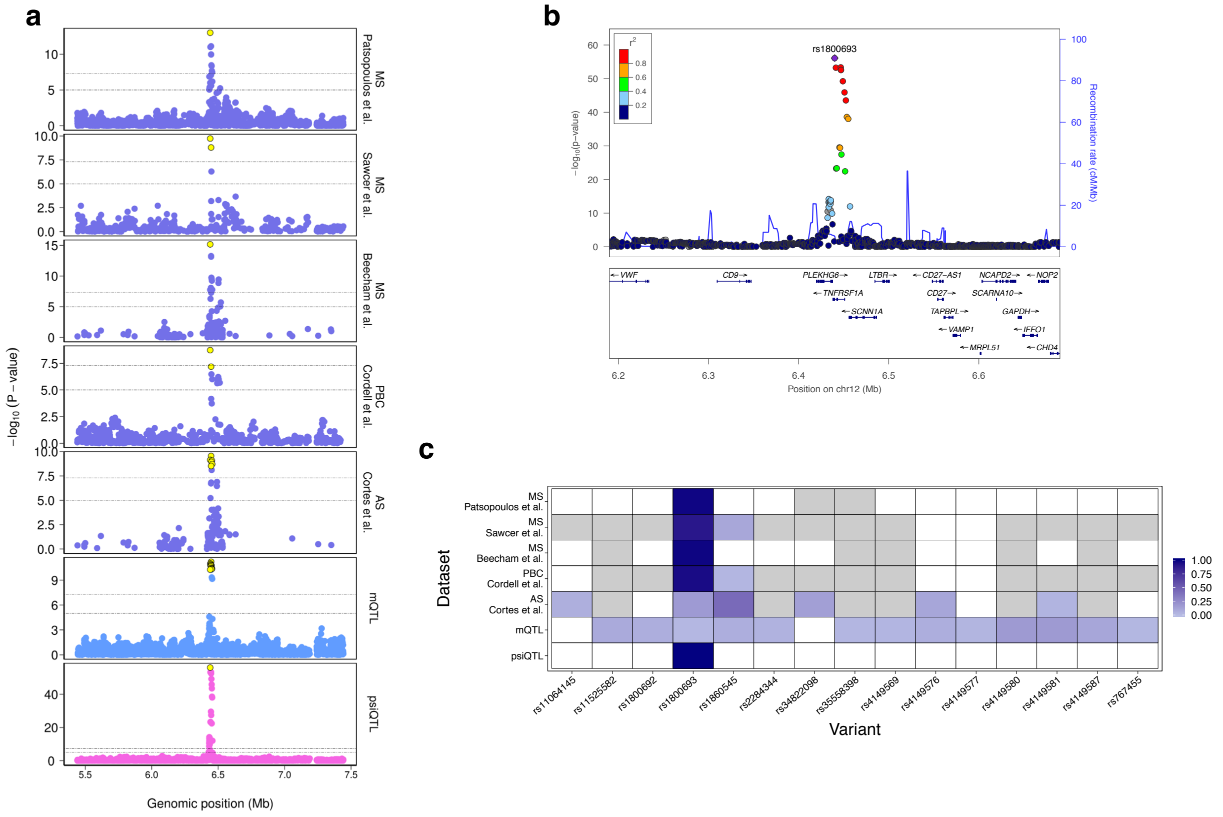


**Supplementary Fig. 19**: *Fine-mapping of the TNFRSF1A locus.* The splicing QTL, rs1800693 (C), results in an increase of a soluble form of *TNFRSF1A* and is a risk allele for MS[3,20,21,25], PBC[26], and highly correlated with rs1860545 ($r^2 = 0.98$), which is a rick allele for AS[27]. **a**, Colocalisation plot of the locus, depicting regional association for the IMD locus for different diseases (blue), and QTLs for psiQTL (pink) and mQTL (lightblue) in monocytes. **b**, A locuszoom plot for the *TNFRSF1A* locus using psiQTL. **c**, Heatmap of posterior probability of variants derived from different credible sets. White boxes indicate the variants are not part of the respective credible sets and grey boxes indicate the variants are actually not included in the respective disease summary statistics. The soluble *TNFRSF1A* inhibits TNF$\alpha$ signaling and mimics the effect of anti-TNF$\alpha$ drugs in MS, where early clinical trials had to be halted due to exacerbation of disease symptoms. The extension of the molecular mechanism underpinning the risk allele from MS to PBC and AS highlights the benefits of the current comprehensive colocalisation effort and argues against the use of anti-TNF$\alpha$ therapies to treat both diseases.

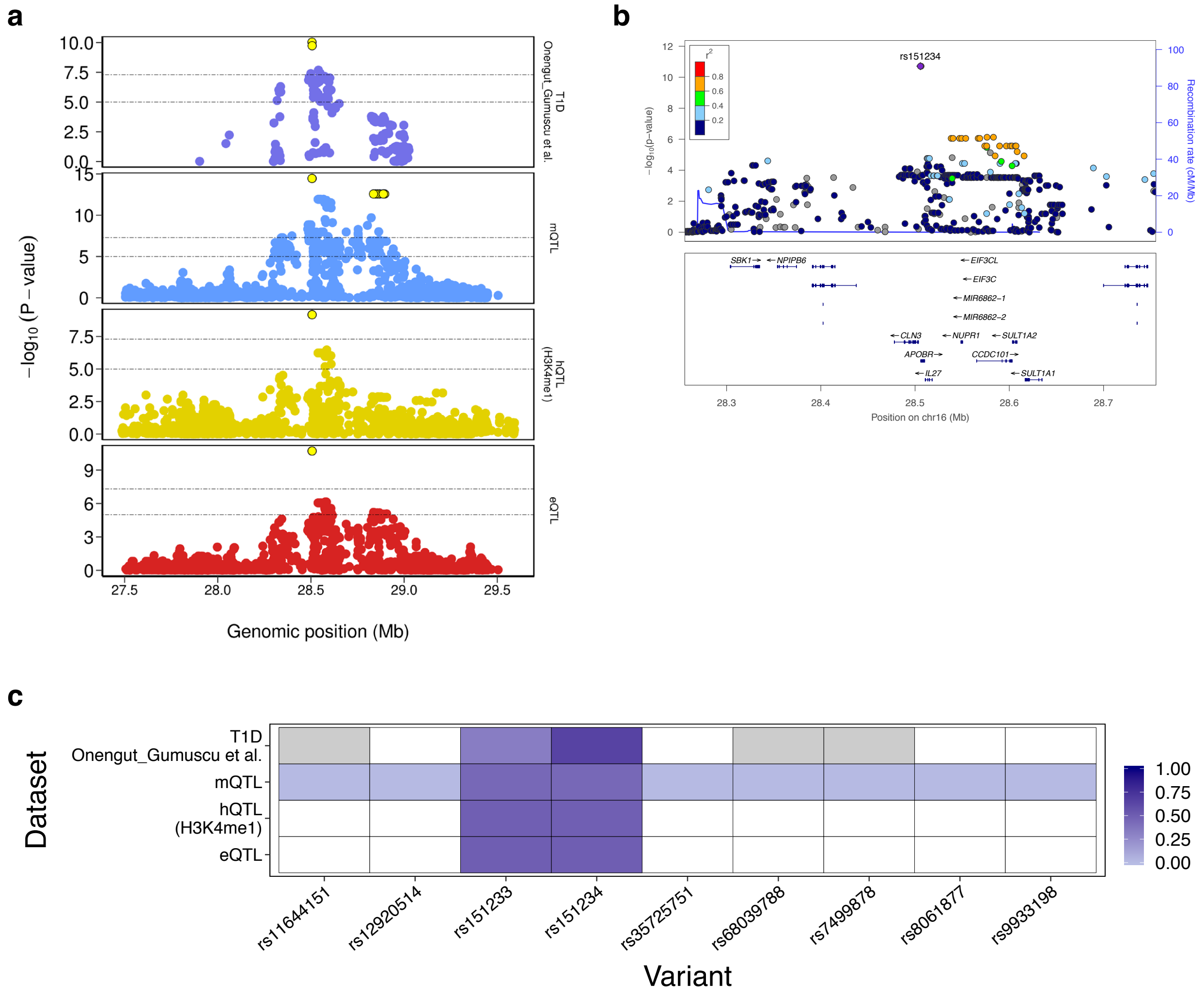


**Supplementary Fig. 20**: *Fine-mapping of the APOBR locus.* The *IL27/CLN3/APOBR* locus is associated with T1D[4] and is colocalised with eQTL (for *APOBR*), hQTLs (H3K4me1), and mQTL in CD4+ T-cells. **a**, Colocalisation plot of the loci, depicting regional association for disease locus (blue), eQTL (red), mQTL (lightblue), hQTL (H3K4me1 in yellow) in CD4+ T-cells. **b**, A locuszoom plot for the *APOBR* locus using eQTL (T-cell) data. **c**, Heatmap of posterior probability of variants derived from different credible sets. White boxes indicate the variants are not part of the respective credible sets and grey boxes indicate the variants are actually not tested in the respective disease datasets. From QTL fine-mapping, rs151233 and rs151234 ($r^2 = 1$ in BLUEPRINT data; both have $PP_{fm} = 0.5$) appear to be more probable causal variants among others. These variants are also part of the IMD credible sets for T1D[4].

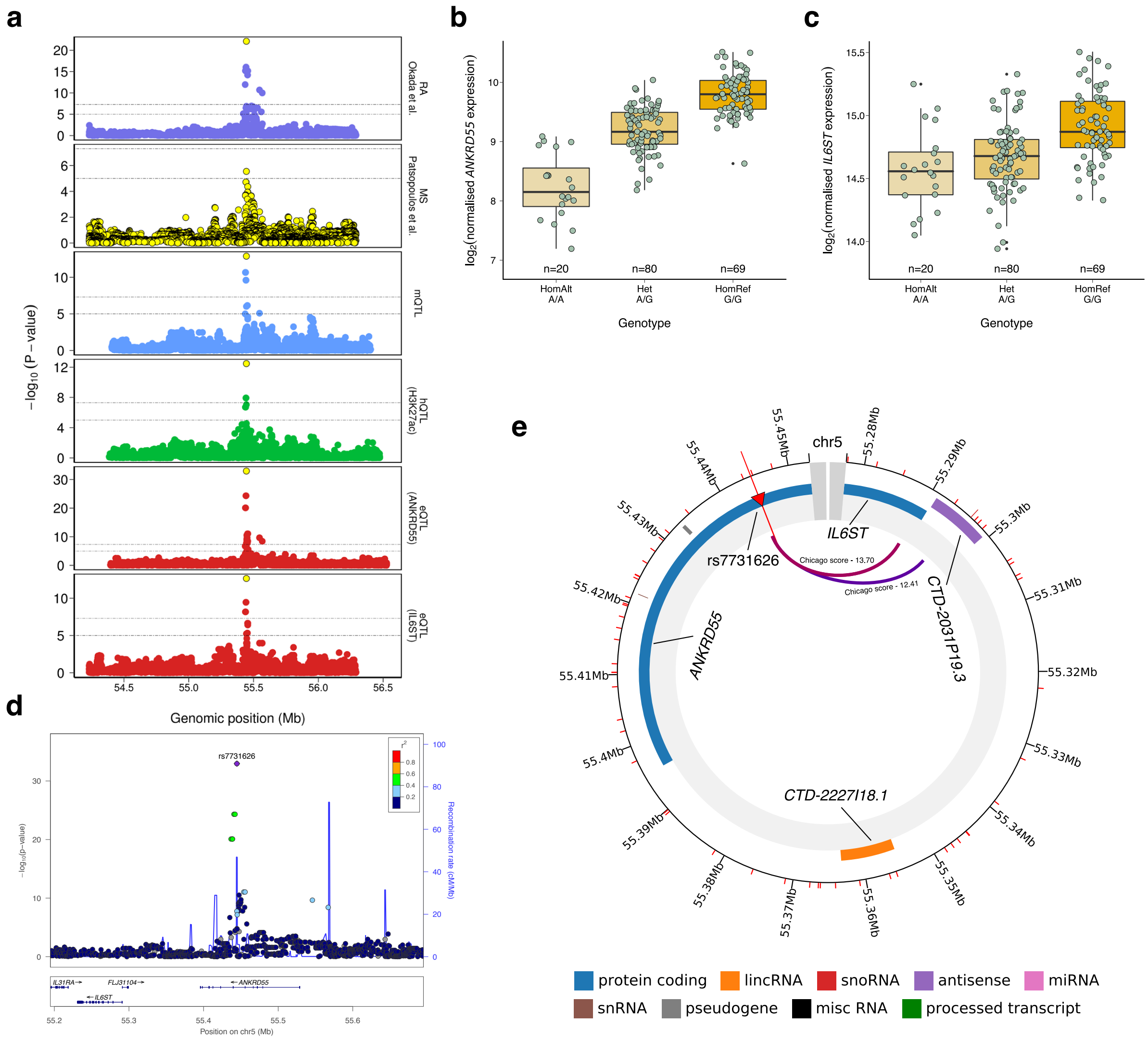


**Supplementary Fig. 21**: *Fine-mapping of the ANKRD55/IL6ST locus.* **a,** We identified an intronic variant, rs7731626 (A) which is a protective allele for MS[3] and RA[10] colocalised with multiple T-cell specific QTLs including eQTLs for two genes, *ANKRD55* and *IL6ST*, H3K27ac and methylation QTLs. Yellow circles represent the variants present in the respective credible sets. The lead variant (rs7731626) appeared to be single fine-mapped variant ($PP_{fm} \sim 1$) for all datasets, except MS, as the signal did not reach genome-wide significant threshold ($P < 5 \times 10^{-8}$) in publicly available summary statistics[3]. Due to lack of power the region could not be fine-mapped using MS summary statistics, however, rs7731626 is the most significant variant and achieved highest posterior probability in the region. **b,c,** The *ANKRD55* and *IL6ST* expression levels affected by rs7731626 in T-cells, respectively. The *ANKRD55* encodes ankyrin repeat domain-containing protein 55, which currently has no known detailed function. *IL6ST* (also known as soluble gp130) encodes a signal transducer involved in multiple cytokine signalling pathways and receptor activation. Blocking IL6 signaling has been hypothesized to limit immune mediated tissue injury in MS, which is supported by preclinical experiments in the mouse EAE model as well as by the treatment of patients suffering neuromyelitis optica with Tocilizumab (IL6 receptor blocker)[28]. **d,** A locuszoom plot using eQTL data with 250kb flanking region surrounding the sentinel SNP (rs7731626). **e,** Using naive CD4+ T-cell PC-HiC data[29], we identified a significant chromatin interaction (Chicago score 13.70) between rs7731626, located within an intron of *ANKRD55*, and the promoter of *IL6ST*, further supporting the eQTL evidence that

**Supplementary Fig. 21**: (continued..) this locus has a pleiotropic mechanistic effect on two genes. This figure was generated by CHiCP server[30]. Our analysis demonstrated that an increase risk in both diseases is correlated with an increase in gene expression of both genes, histone activity and a reduction in DNA methylation, suggesting an immune role in the function of both genes and in their involvement in immune-mediated disease risk. The colocalisation of this locus with *ANKRD55* expression has previously been identified but not colocalised with *IL6ST*[31]. Using more variant-dense data for RA[10], we identified rs7731626 was the single causal variant in the credible set for both genes and RA, but was previously missed from lower-powered studies[32,33]. This suggests rs7731626 was the most likely causal variant and has a role on two genes.
